# TransCell: In silico characterization of genomic landscape and cellular responses from gene expressions through a two-step deep transfer learning

**DOI:** 10.1101/2022.05.19.492662

**Authors:** Shan-Ju Yeh, Shreya Paithankar, Ruoqiao Chen, Jing Xing, Mengying Sun, Ke Liu, Jiayu Zhou, Bin Chen

## Abstract

Gene expression profiling of new or modified cell lines becomes routine today; however, obtaining comprehensive molecular characterization and cellular responses for a variety of cell lines, including those derived from underrepresented groups, is not trivial when resources are minimal. Using gene expression to predict other measurements has been actively explored; however, systematic investigation of its predictive power in various measurements has not been well studied. We evaluate commonly used machine learning methods and present TransCell, a two-step deep transfer learning framework that utilizes the knowledge derived from pan-cancer tumor samples to predict molecular features and responses. Among these models, TransCell has the best performance in predicting metabolite, gene effect score (or genetic dependency), and drug sensitivity, and has comparable performance in predicting mutation, copy number variation, and protein expression. Notably, TransCell improved the performance by over 50% in drug sensitivity prediction and achieved a correlation of 0.7 in gene effect score prediction. Furthermore, predicted drug sensitivities revealed potential repurposing candidates for new 100 pediatric cancer cell lines, and predicted gene effect scores reflected BRAF resistance in melanoma cell lines. Together, we investigate the predictive power of gene expression in six molecular measurement types and develop a web portal (http://apps.octad.org/transcell/) that enables the prediction of 352,000 genomic and cellular response features solely from gene expression profiles.

**Key Points:**

- We provide a systematic investigation on evaluating the predictive power of gene expression in six molecular measurement types including protein expression, copy number variation, mutation, metabolite, gene effect score, and drug sensitivity.
- TransCell took advantage of the transfer learning technique, showing how to learn knowledge from the source tumors, and transfer learned weight initializations to the downstream tasks in cell lines.
- Compared to the baseline methods, TransCell outperformed in metabolite, gene effect score, and drug sensitivity predictions.
- Two cases studies demonstrate that TransCell could identify new repurposing candidates for pediatric cancer cell lines as well as capture the differences of genetic dependencies in melanoma resistant cell lines.

## INTRODUCTION

*In vitro* cell lines are widely used in biomedical research and drug discovery [1-3]. New cell lines are emerging rapidly thanks to the advances in the cell culture of individual tumors, and existing cell lines are frequently modified (e.g., drug treatment, CRISPR knockout) to delineate underlying biology. Molecular profiling of these cell lines and large-scale characterization of their responses to perturbagens allows the discovery of cancer driver genes, therapeutic candidates, and biomarkers. Cell line gene expression profiling becomes routine today; however, it remains costly to generate other data types for individual cell lines where resources are often limited. Consequently, those data derived from underrepresented groups (e.g., age: pediatric, race: African American, American Indian) are scarce. The recent efforts in large-scale profiling of common cancer cell lines provide a rigorous framework for studying other cancers. Outstanding examples include DepMap (a collection of projects such as CCLE, PRISM, and Achilles), where transcriptomic, proteomic, metabolomic, and genomic features of over 1000 cancer cell lines, as well as genetic screenings and drug screenings in these cell lines, were recently profiled [4]. RNA, as an intermediate product between DNA and protein, is known to encode information of other measurements (even including potential responses to external agents). If a computational model could decode the information solely based on gene expression, then *in silico* characterization of genomic landscape and cellular responses from gene expressions for new cell lines is within reach.

Using gene expression to predict the measurement of individual feature types has been actively explored. To achieve optimal performance, genomics and other omics data are frequently borrowed in the prediction of drug sensitivity [5-9], drug synergy [10-12], and gene essentiality [13-15]. Gene expression, mutation and copy number variation (CNV) were also combined in drug response prediction [16]. Moreover, in drug-related tasks, irregular or non-Euclidean drug structures could be embedded through convolution neural networks (CNN) [8, 17, 18] and graph representations [19-21] to aid the modeling. Nevertheless, gene expression consistently shows the most informative in drug response prediction [22, 23], whereas mutation and CNV profiles contribute little to improve the accuracy [24]. Although the integration of multiple omics profiles could improve the learning performance, its application is limited in practice where only gene expression data is often accessible. The aforementioned prior studies inspired us to perform a systematic investigation of the power of using gene expression in the prediction of various measurement types available in the recent DepMap collection.

In this study, using DepMap data, we comprehensively evaluate commonly used feature selection and baseline machine learning methods in the prediction of protein expression, CNV, mutation, metabolite, gene effect score (a measurement of genetic dependency from CRISPR screening), and drug sensitivity (Figure 1a-b). To address the big p (features) and little n (samples) problem, we further develop TransCell, a deep transfer framework that leverages a pre-training model from large TCGA pan-cancer patient profiles to gain better weight initializations. Comparing to train DNN directly, we demonstrate the superiority of TransCell, which could overcome premature convergence and overfitting under a relatively small sample size. TransCell outperforms baseline machine learning methods in metabolite, gene effect score, and drug sensitivity prediction. Meanwhile, TransCell is readily accessible and convenient to use in the prediction of six molecular measurement types solely based on gene expression through the web portal (http://apps.octad.org/transcell/)

**Figure 1.**
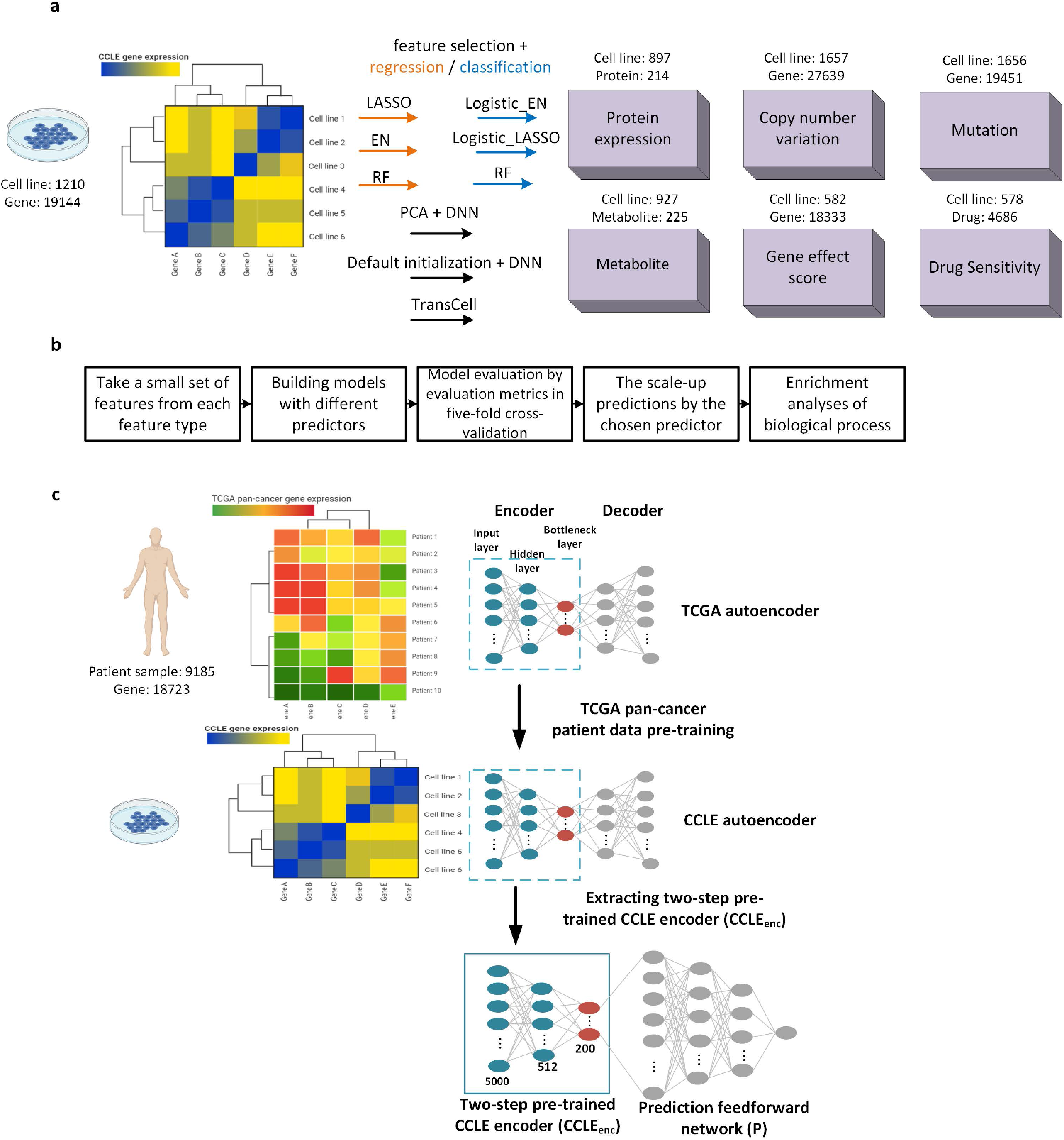
An overview of research design. **a**. Prediction of measurements in six types based on cancer cell line gene expression data. The number of cell lines varies across data types. LASSO: Least Absolute Shrinkage and Selection Operator, EN: Elastic Net, RF: Random Forest, PCA: Principal Component Analysis, DNN: Deep Neural Network. **b**. Model evaluation process. Due to the high demand for computation power, we started with a small set of measurements for each type and then scaled up to a larger set. **c**. Schematic of TransCell. The top 5000 features sharing similar distribution between CCLE and TCGA were first selected, followed by the creation of an autoencoder using TCGA pan-cancer tumor transcriptomics. The parameters of the TCGA encoder were then transferred to the second CCLE autoencoder for weight initializations. Afterward, a two-step pre-trained CCLE encoder was extracted and linked to a prediction feedforward network. Parameters were tuned automatically (see Methods). Note that one model is built for each molecular measurement.

## MATERIAL AND METHODS

### Dataset

The cancer cell line log2 transformed TPM gene expression matrix for protein-coding genes (processed by RSEM) including 1210 cell lines and 19144 genes was downloaded from the DepMap portal (http://www.depmap.org) [4]. The TCGA pan-cancer RSEM-processed gene expression matrix including 9185 patient samples and 18,723 genes was downloaded from OCTAD [25]. The measurements, including protein expression, CNV, metabolite, gene effect score, drug sensitivity, and mutation data, were downloaded from the DepMap portal [26-28] (Figure 1a). The DepMap 20Q1 serving as a prospective dataset was downloaded from this link (https://figshare.com/articles/dataset/DepMap_20Q1_Public/11791698). For new RNASeq profiles, we downloaded the raw sequences from SRA and used the OCTAD pipeline to compute TPM [25].

### Transfer learning between TCGA and CCLE

TransCell was developed to predict protein expression, CNV, metabolite, gene effect score, drug sensitivity, and mutation data using gene expression features taken from CCLE. As shown in Figure 1c, TransCell is composed of two networks: i) a two-step pre-trained CCLE encoder (CCLE_enc_) and ii) a prediction feedforward network (P). The first component is an encoder extracted from the second autoencoder trained by CCLE, transforming higher-order features of CCLE gene expression into a lower-dimensional representation. The encoded representation is then linked to P, resulting in a fully connected network for training. We adopted a parameter-based transfer learning, where the weights learned from the source domain are transferred to the target domain for weight initializations, followed by finetuning all layers. One major concern of transfer learning is negative transfer, meaning that the source domain needs to resemble the target domain; otherwise, the attempt to transfer knowledge from the source can have a negative impact on the target learner and thus can even lower the performance [29]. Considering feature distribution within source domain (TCGA) and target domain (CCLE) for minimizing the effects of negative transfer, we performed a two-sample Kolmogorov-Smirnov test for each gene feature to identify genes with a similar or identical distribution between TCGA and CCLE. According to adjusted p values, we chose the top 5000 genes to be our features for training TransCell.

### Two-step pre-training of cancer cell line encoder (CCLE_enc_)

Autoencoder, a non-linear unsupervised learning, comprises a symmetric pair of an encoder and a decoder. Given a dataset *X* (*X* = {x_1_,…, x_*n*_}) with *n* samples and *m* features, the encoder is a function *f* that maps the input *X* to a lower-dimensional representation, which could be described as below:

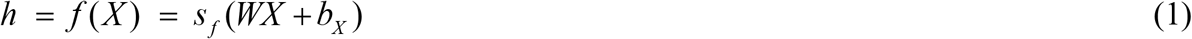

where *h* represents the hidden representation of *X, s* _*f*_ stands for LeakyReLU, a nonlinear activation function, making autoencoder perform a nonlinear projection [30]. The encoder is parameterized by a weight matrix *W* and a bias vector *b*_*X*_. Moreover, the decoder function *g* maps the hidden representation *h* back to a reconstruction 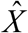. The process is formulated in the following:

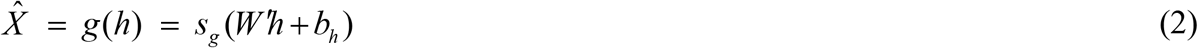

where *s*_*g*_ is an identity. The decoder’s parameters comprise a weight matrix *W* ′ and a bias vector *b*_*h*_. The training process of an autoencoder aims to find a parameter set *θ*= {*W, W* ′, *b*_*X*_, *b*_*h*_} that minimizes the reconstruction loss on the given data dataset *X*. The objective function is given as

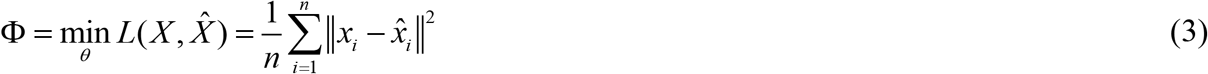

By minimizing the loss between the original input *X* and reconstruction 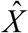 in the equation (3), an autoencoder reduces the dimension of complex input features and produces a compressed representation at the bottleneck layer, the layer between encoder and decoder. We pre-trained an encoder using a large TCGA set and transferred its weights to the CCLE autoencoder as its weight initializations and then extracted two-step pre-trained CCLE_enc_ to link to P. We trained the autoencoder using 90% of CCLE data together with 10% as the validation set. The optimal hyperparameters were searched by Keras Tuner (https://keras-team.github.io/keras-tuner/) based on the loss of the validation set. As a result, for the architecture of encoder, the input layer had 5000 neurons, followed by one hidden layer with 512 neurons, and a bottleneck layer with 200 neurons. The activation function was set as LeakyReLU (alpha=0.1) for each layer in the autoencoder. The complete algorithm for two-step pre-training CCLE_enc_ is given in Figure 2.

**Figure 2.**
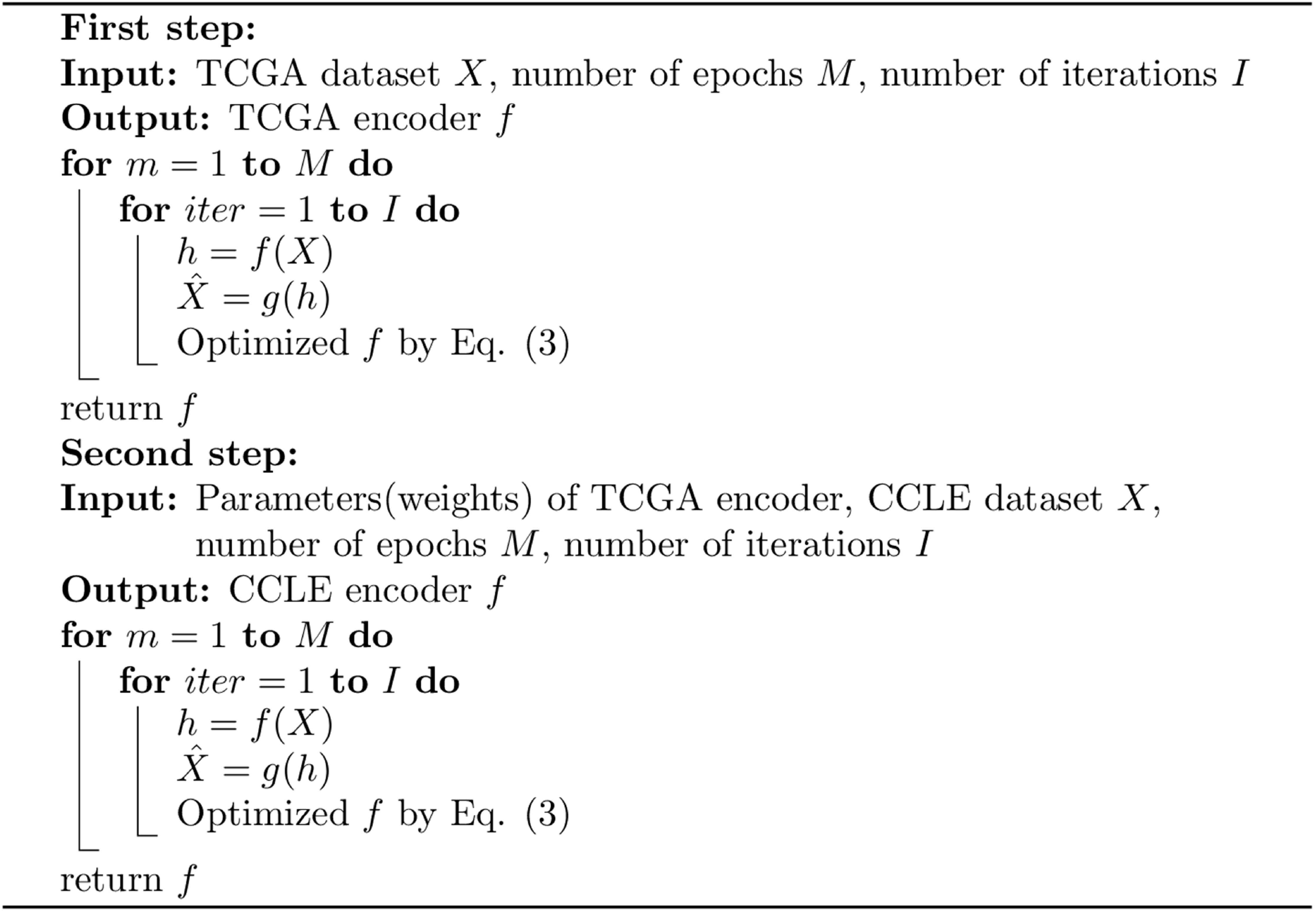
Pseudocode of two-step pre-training CCLE encoder.

### Prediction network

Following the two-step pre-trained CCLE_enc_ is P, a 4-layer feedforward neural network, including the first layer merging output neurons of the two-step pre-trained CCLE_enc_, two fully connected layers, and one output layer generating predicted values. The architecture of two-step pre-trained CCLE_enc_ was fixed. Its initialized parameters came from two-step pre-training and the weights would be updated during the training process. P used default initialization. To enable various partitions of data as required by the different prediction scenarios and to obtain robust evaluation metrics including mean squared error (MSE), root mean squared error (RMSE), and Spearman rank correlation, we used the average of five-fold cross-validation results for model evaluation. For each time, four folds of data randomly split from the whole data were used together with 10% validation set and one fold of data was used as a testing set. The architectures for the measurement types are detailed in Supplementary Materials.

### Feature Selection methods

TransCell obtained compressed CCLE representation from the encoder. Except for using encoder capturing essential feature information, we investigated the common feature selection and dimension reduction methods, including least absolute shrinkage and selection operator (LASSO), elastic net (EN), random forest (RF), and principal component analysis (PCA) (Supplementary Materials).

### Comparison to other model designs

One primary reason to evaluate feature selection and dimension reduction methods, as well as predictors, is to choose the best available model and estimate how well a given model is likely to perform in making predictions for each measurement type. The performance of TransCell was compared to three baseline machining learning methods, including LASSO, EN, and RF, and two DNN designs. Many existing algorithms were developed for individual tasks, thus were not included for a back-to-back comparison, but we investigated some individually. The LASSO predictor relied on LASSO to perform feature selection and chose all the features with non-zero coefficient to be the essential features that were later selected to build a LASSO model. The EN predictor included an elastic net to identify all features with non-zero coefficients as essential features, followed by an elastic net model. The RF predictor used its internal feature selection method to build the model with feature importance > 10^-4^ as essential features.

We designed two similar DNN architectures for comparisons, with one replacing two-step pre-trained CCLE_enc_ with PCA (using 200 principal components instead of the 200 compressed autoencoder features) and another training with default initializations without transferred parameters. To conduct a performance comparison in mutation prediction, a logistic regression model was applied for both feature selection and classification. Note that we compare the logistic regression model with *l*_1_ regularization (Logistic_LASSO) and the logistic regression model with the combination of *l*_1_ and *l*_2_ regularizations (Logistic_EN), two different ways of feature selection in logistic regression. To reduce the searching time of hyperparameters, for each type of predictor in the prediction for each measurement type, we derived hyperparameters based on one model and applied them to all models. The optimized hyperparameters are found by scikit-learn’s GridSearchCV for LASSO, EN, RF, Logistic_LASSO, and Logistic_EN. During cross-validation tuning, we used MSE as a scoring metric for the best hyperparameters search. The actual values we used for feature selection and prediction model in each type of predictor are shown in Supplementary Table 1. The computing time for each predictor to obtain an average of five-fold cross-validation results is around five minutes.

### Model evaluation

MSE and RMSE are commonly used model evaluation metrics for measuring model performance. Both MSE and RMSE are sensitive to outliers, meaning that they give more weight to larger differences. Because of giving higher weight to unfavorable conditions, they usually are good at revealing model performance differences [31]. In order to make further intuitive assessments, we also computed Spearman rank correlation between predicted and actual values. It is noted that RMSE is not only interpretable in terms of measurement units but also a better measurement of model fit than a correlation coefficient [32]. Therefore, we used RMSE to derive well-predicted and poorly predicted features, upon which gene or metabolite set enrichment analyses of biological processes were performed. To avoid overfitting and biased results, the evaluation metrics were computed based on the average of five-fold cross-validation results. In the five-fold cross-validation, hyperparameters were fixed, each of the k fold is given an opportunity to be used as a hold-out test set while all other k-1 folds are used as a training set. This procedure would repeat for k times iteratively. Therefore, we could compute the mean of evaluation metrics.

### Pediatric cancer sensitivity prediction

The gene expression profiles of pediatric cancer cell lines were downloaded from [33] and then fed into trained models. Drug sensitivity data were collected from PubChem Bioassay (RDES: AID1259252, TC32: AID1259256, EW8: AID1259255). PubChem ID Exchange service was used to convert SMILES in DepMap to PubChem CID, which was later used to map drug sensitivity. PUBCHEM_ACTIVITY_OUTCOME was compared with predicted drug sensitivity. We considered predicted score < -2 as active compounds (a few other thresholds were explored as well). Fisher exact test was applied to identify selective compounds for each cancer. R packages umap and ComplexHeatmap were used to visualize cell lines and drug sensitivity.

## RESULTS

We developed a pipeline to systematically evaluate models in the prediction of protein expression, CNV, mutation, metabolite, gene effect score, and drug sensitivity using gene expression data from CCLE (Figure 1a-b). The gene expression matrix consists of 1210 cell lines and 19,144 genes, while the six measurement types have a varying number of cell lines and measurements (Figure 1a). All these measurements are continuous except mutation, a binary measurement. For each measurement type, we started with a small set of measurements for the following predictors: LASSO, EN, RF, PCA features + DNN, and DNN + default initialization (Methods). For mutation data, we considered Logistic_LASSO and Logistic_EN. Each model comprises feature selection and regression/classification (Figure 1a). For evaluation metrics, we employed the two most frequently used criteria: MSE and RMSE [34]. In addition to model errors provided by MSE and RMSE, Spearman rank correlation was added to measure the model agreement between predicted values and real values. Five-fold cross-validation was applied to each predictor. The best available predictor, chosen based on the evaluation metrics including MSE and RMSE, and Spearman rank correlation, was then adopted to perform the scale-up prediction. Due to the constraint of computation power, we randomly selected 2000 measurements for each type if the total measurements exceed 2000. Moreover, gene enrichment analyses of the well- and poorly predicted measurements were performed using the R package clusterProfiler [35] (Figure 1b).

We further developed TransCell, a framework for predicting molecular measurements by utilizing the transfer learning technique. TransCell first builds an autoencoder using a larger dataset comprising about 10,000 gene expression profiles from TCGA. The parameters of the resulting encoder are used to initialize the autoencoder built from CCLE gene expression profiles. In light of the difference between tumors and cell lines, TransCell only encodes the genes sharing a similar distribution between the two sets. Subsequently, a two-step pre-trained CCLE_enc_ is extracted and then linked to a feedforward prediction network (P).

### Metabolite prediction

Metabolomics analyses facilitate the identification of biomarkers and improve the understanding of biological pathways in health and metabolic diseases. Among the six molecular types in DepMap, metabolomics is the last one becoming publicly available [26]. Its prediction using gene expression is barely explored, thus we first start with metabolite prediction. In order to quickly sense which predictor has the best performance, we randomly chose 20 metabolites to build a model for each. After removing missing values, 915 cell lines were left for prediction. A model is built for each metabolite individually. Among the predictions of 20 metabolites, TransCell reached the mean of MSE and RMSE 0.064 (95% confidence interval (CI): [0.051-0.076]) and 0.228 (95% CI: [0.204-0.252]), respectively, which are lower than LASSO, EN, and RF (Figure 3a-b). Furthermore, TransCell outperformed DNNs with PCA or default initialization. The mean of Spearman rank correlations computed between true values and predicted values among 20 models is 0.744 (95% CI: [0.724-0.764]) in TransCell, which is substantially higher than other predictors (Figure 3c). The evaluation metrics and Spearman rank correlation suggest that TransCell has the best performance in metabolite prediction. Therefore, we adopted TransCell to predict all 225 metabolites. In the scale-up prediction, the mean of MSE, RMSE, and Spearman rank correlation is 0.072 (95% CI: [0.064-0.080]), 0.237 (95% CI: [0.226-0.247]), and 0.746 (95% CI: [0.740-0.753]), respectively (Figure 3d-f). Among those metabolites, the top five well-predicted metabolites are C34:1 PC, C36:2 PC, C16:1 SM, C34:2 PC, and C38:4 PC. The top five poorly predicted metabolites are acetylcholine, taurochenodeoxycholate, methylnicotinamide, hypoxanthine, and palmitoylcarnitine.

**Figure 3.**
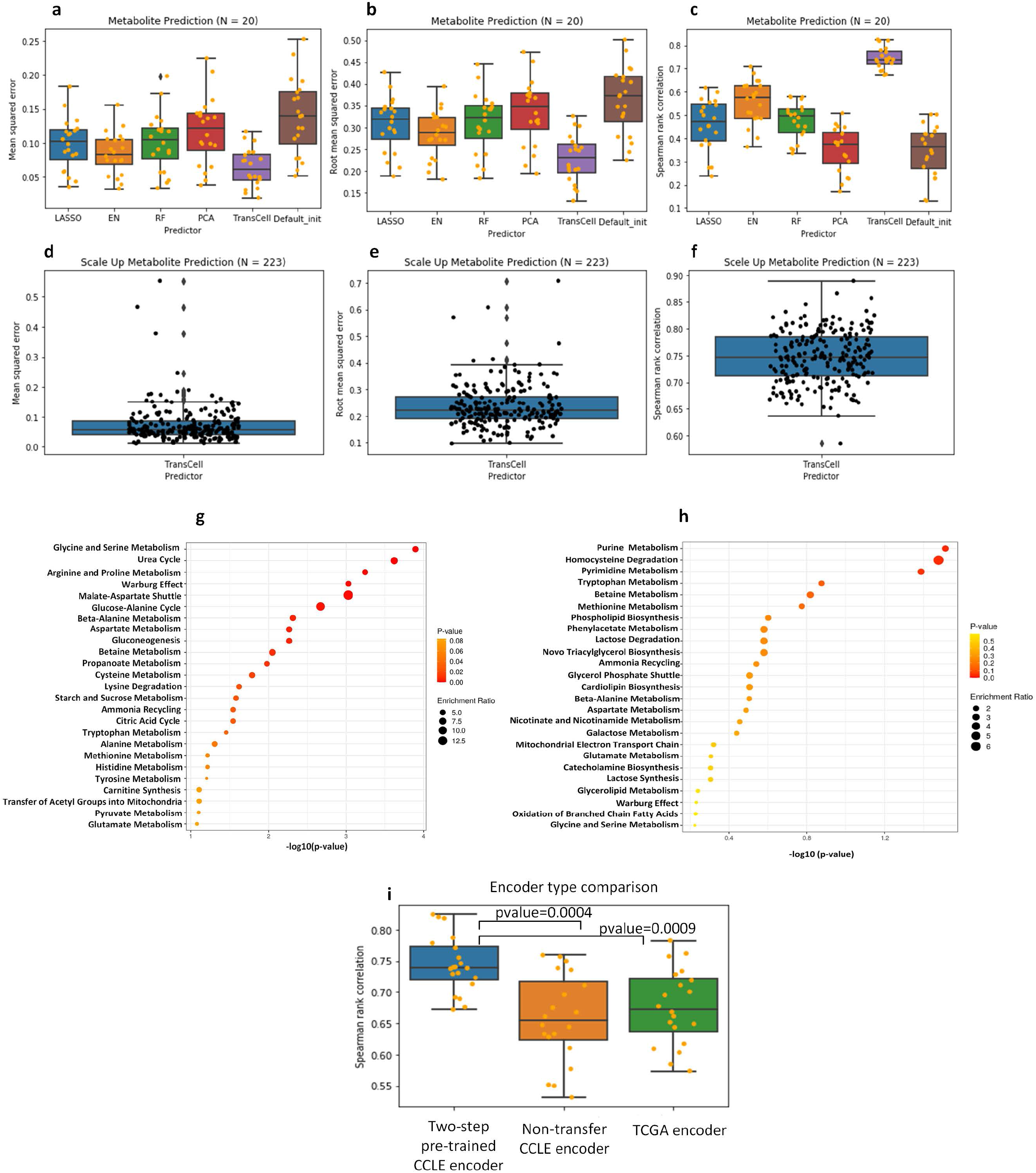
Metabolite predictions. **a-c**. The boxplots of (a) MSE, (b) RMSE, and (c) Spearman rank correlation for different predictors, including LASSO, EN, RF, TransCell, and two DNN designs with PCA and default initializations in 20 metabolite prediction models. **d-f**. The boxplots of (d) MSE, (e) RMSE, and (f) Spearman rank correlation for TransCell in all 223 scale-up metabolite prediction models. **g**. The dot plot of metabolite set enrichment analyses for metabolites with RMSE lower than the first quartile (well-predicted metabolites) in all 223 scale-up metabolite predictions. **h**. The dot plot of metabolite set enrichment analyses for metabolites with RMSE greater than the third quartile (poorly predicted metabolites) in all 223 scale-up metabolite predictions. **i**. Spearman rank correlation for the TransCell architecture using two-step pre-trained CCLE encoder, non-transfer CCLE encoder, and TCGA encoder.

To investigate well- and poorly predicted metabolites, we derived those with RMSE lower than the first quartile as well-predicted metabolites, and those with RMSE greater than the third quartile as poorly predicted metabolites. After that, we performed metabolite enrichment analyses by MetaboAnalyst [36]. The enrichment showed that the top three pathways related to well-predicted metabolites were glycine and serine metabolism, urea cycle, and arginine and proline metabolism (Figure 3g). The pathways involved in purine metabolism, homocysteine degradation, and pyrimidine metabolism were related to poorly predicted metabolites (Figure 3h).

To demonstrate the efficacy of the two-step pre-trained CCLE_enc_, we explored another two types of encoders. The first type of encoder, CCLE_enc_, was trained by CCLE without transferred encoder parameters learned from TCGA. The second type of encoder, TCGA encoder (TCGA_enc_), was trained by TCGA. We repeated the analysis using the results from each new encoder. We found that TransCell using the two-step pre-trained CCLE_enc_ has the best performance (P < 0.05 using Wilcoxon rank-sum test, Figure 3i), indicating that TransCell offers better parameter weight initializations.

### Gene effect score prediction

Similar to metabolite prediction, we first randomly chose 20 genes to build gene effect score prediction models. After removing missing values, 578 common cell lines between CCLE and DepMap were left. Among 20 gene effect score models built by TransCell, the mean of MSE and RMSE is 0.015 (95% CI: [0.010-0.020]) and 0.105 (95% CI: [0.091-0.119]), respectively, and the mean of Spearman rank correlation is 0.689 (95% CI: [0.672-0.707]) (Figure 4a-c). TransCell has the lowest MSE and RMSE compared to LASSO, EN, RF, and two DNN designs with PCA and default initializations, and holds the lowest variation and the highest Spearman rank correlation with significant p-values (P < 0.05, Student’s t-test), suggesting the superiority of TransCell in gene effect score prediction. Therefore, we used TransCell to predict 2000 randomly selected genes. The mean of MSE, RMSE, and Spearman rank correlation within these 2000 models is 0.012 (95% CI: [0.011-0.012]), 0.095 (95% CI: [0.094-0.096]), and 0.676 (95% CI: [0.675-0.678]) (Figure 4d-f), respectively, corroborating with the small-scale test.

**Figure 4.**
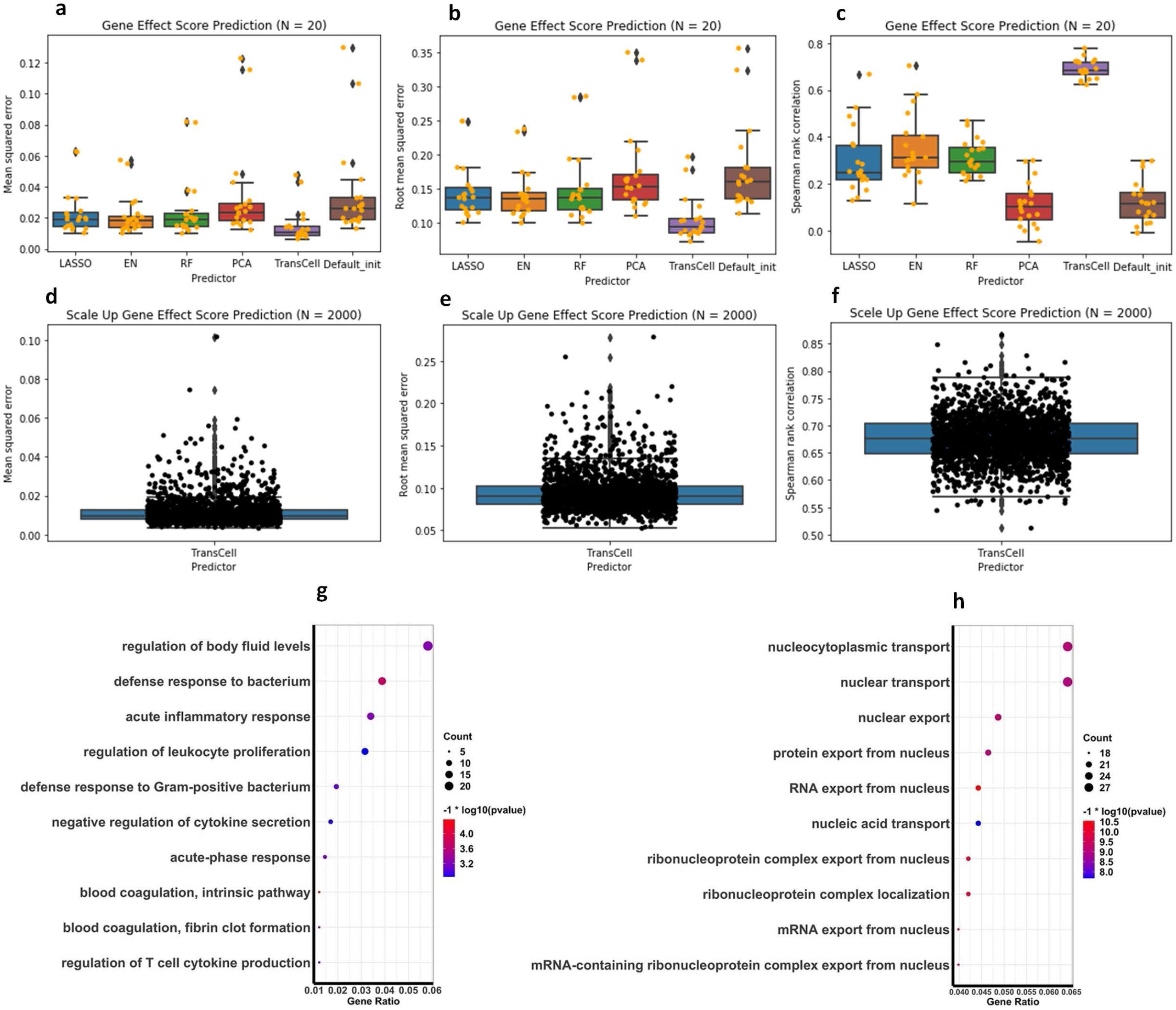
Gene effect score predictions. **a-c**. The boxplots of (a) MSE, (b) RMSE, and (c) Spearman rank correlation for different predictors, including LASSO, EN, RF, TransCell, and the two DNN designs with PCA and default initializations in 20 gene effect score prediction models. **d-f**. The boxplots of (d) MSE, (e) RMSE, and (f) Spearman rank for TransCell in 2000 scale-up gene effect score prediction models. **g-h**. The dot plots of gene enrichment analysis of biological processes for genes with RMSE (g) lower than the first quartile (well-predicted genes) and (h) higher than the third quartile (poorly predicted genes) in the scale-up gene effect score predictions.

Based on RMSE distribution, we further examined well- and poorly predicted genes (poorly: greater than the third quartile, well: smaller than the first quartile). Gene enrichment analyses found the top three pathways related to well-predicted genes were (1) regulation of body fluid levels, (2) defense response to bacterium, and (3) acute inflammatory response, and those related to poorly predicted genes are (1) nucleocytoplasmic transport, (2) nuclear transport, and (3) nuclear export (Figure 4g-h). The poor performance of transporter genes might suggest the irrelevance of gene expression in transporter function.

### Drug sensitivity prediction

In order to investigate which predictor has the best performance for drug sensitivity prediction, 20 compounds were chosen randomly to build models. TransCell again outperformed others with the mean of MSE, RMSE and Spearman correlation 0.207 (95% CI: [0.094-0.320]), 0.375 (95% CI: [0.295-0.455]) and 0.653 (95% CI: [0.637-0.669]), respectively (Figure 5a-c). Hence, using TransCell, we performed scale-up drug sensitivity prediction in 2000 compounds selected randomly. The mean of MSE, RMSE, and Spearman rank correlation within these 2000 models is 0.171 (95% CI: [0.162-0.180]), 0.341 (95% CI: [0.334-0.348]), and 0.657 (95% CI: [0.655-0.659]), respectively (Figure 5d-f). One recent study using RF achieved the Pearson correlation of 0.389, which is close to our observation (Figure 5c) [37]. TransCell improved the correlation by over 50%. We next selected well- and poorly predicted compounds and mapped them to their putative targets through the drug-target mappings provided in DepMap. Pathway enrichment analyses of these targets revealed that sensitivities of the compounds involved in the regulation of membrane potential, regulation of ion transmembrane transport, and divalent inorganic cation transport were accurately predicted, while those targeting calcium ion transport, divalent metal ion transport, and divalent inorganic cation transport were poorly predicted (Figure 5g-h).

**Figure 5.**
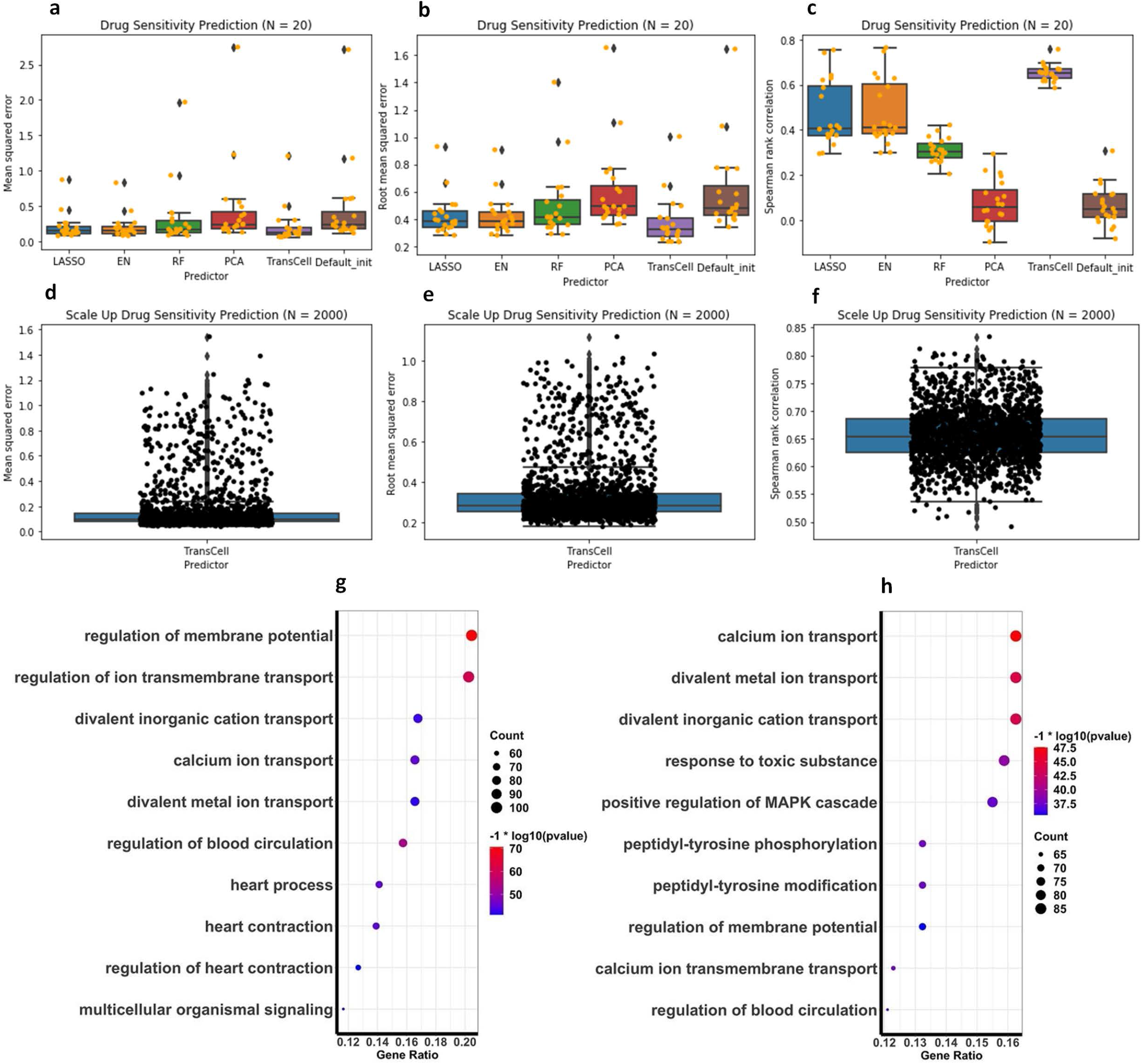
Sensitivity predictions. **a-c**. The boxplots of (a) MSE, (b) RMSE, and (c) Spearman rank correlation for different predictors, including LASSO, EN, RF, TransCell, and two DNN designs with PCA and default initializations in 20 drug sensitivity prediction models. **d-f**. The boxplots of (d) MSE, (e) RMSE, and (f) Spearman rank for TransCell in 2000 scale-up drug sensitivity prediction models. **g-h**. The dot plots of gene enrichment analysis of biological processes for target genes of compounds with RMSE (g) lower than the first quartile (well-predicted compounds) in the scale-up drug sensitivity prediction and (h) higher than the third quartile (poorly predicted compounds) in the scale-up drug sensitivity predictions.

### Protein, copy number variation, and mutation prediction

Unlike the previous three measurement types where TransCell performs the best, protein prediction tends to favor linear-based models like EN, which has the best performance with a Spearman rank correlation of 0.794 (95% CI: [0.745-0.843]) (Figure 6a, Figure S1a-c). The performance of TransCell is comparable, with no significant difference detected (Spearman rank correlation difference, t-test p-value: 0.153). While scaling up to all 214 proteins using EN, the performance is stable (Spearman correlation 0.792 (95% CI: [0.777-0.806])) (Figure S1d-f). Diving into individual proteins, we observed that the proteins related to apoptotic signaling pathway, regulation of protein complex assembly, and epithelial cell proliferation could be easily predicted. In contrast, those related to serine/threonine kinase activity, response to toxic substance, and response to nutrient levels are less likely to be inferred from gene expression (Figure S1g-h).

**Figure 6.**
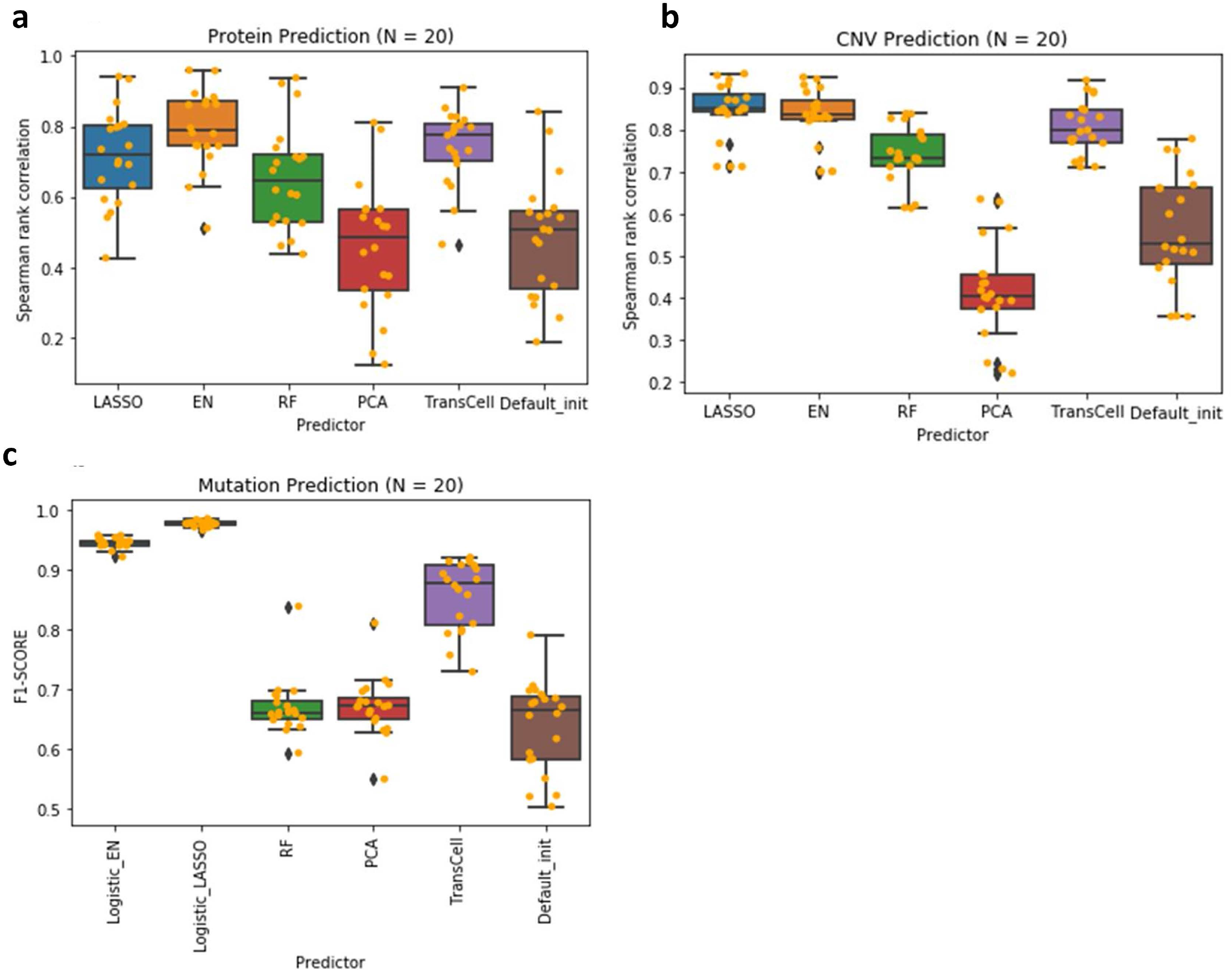
Protein, copy number variation, and mutation prediction. **a-b**. Spearman rank correlation was used to measure the performance in protein prediction (a) and CNV prediction (b). **c**. Mutation prediction was considered as a classification problem; thus F1-score was chosen to evaluate the performance. Full prediction performance is in Supplementary Figure 1-3.

Similar to protein expression, CNV can be easily predicted by linear models. LASSO stood out, with the mean of MSE, RMSE, and Spearman rank correlation 0.009 (95% CI: [0.006-0.012]), 0.087 (95% CI: [0.070-0.103]), and 0.844 (95% CI: [0.814-0.874]), respectively (Figure 6b, Figure S2a-c). TransCell and EN have comparable performance with LASSO (p-value > 0.01). Subsequent modeling of 2000 genes using LASSO suggests the consistent power of predicting CNV using gene expression alone (Spearman correlation 0.874 (95% CI: [0.872-0.876])) (Figure S2d-f). For the handful of poorly predicted genes, no prominent patterns were observed (Figure S2g-h).

In mutation prediction, we used AUC and F1 score to evaluate the classification of mutation status (mutated/wild). Logistic_LASSO and Logistic_EN have the best performance, followed by TransCell. Among 20 mutation prediction models built by Logistic_LASSO, the mean of AUC and F1 score is 0.996 (95% CI: [0.995-0.997]) and 0.978 (95% CI: [0.976-0.980]), respectively, suggesting that mutation could be precisely predicted by gene expression (Figure 6c, Figure S3a-b). While scaling up to 2000 mutations using Logistic_LASSO, nearly all the mutations could be well-predicted shown in Figure S3c-d (AUC: 0.990 (95% CI: [0.9896-0.9903]) and F1: 0.971 (95% CI: [0.9703-0.9709])). In order to explore poorly predicted genes in the scale-up mutation prediction, we considered genes with AUC lower than the first quartile as poorly predicted. Afterward, we performed gene enrichment analyses of the biological process for those genes. The top three pathways ranked by gene ratio were (1) cell morphogenesis involved in neuron differentiation, (2) synapse organization, and (3) axon development (Figure S3e).

### Model evaluation using external and prospective datasets

Having shown the feasibility of using gene expression to predict other molecular measurements, mainly using DepMap, we next sought external validation to verify the robustness of the models. We realized the datasets with both RNA-Seq and other measurements are surprisingly scarce; we could only explore a few datasets. First, we analyzed proteomic data in CellMinerCDB (discover.nci.nih.gov/cellminercdb). The protein array data in CellMinerCDB were taken from MDACC CLP (https://tcpaportal.org/mclp), and the RNA-Seq data were based on the NCI60-cell lines. There are 60 cell lines and 36 common proteins between CellMinerCDB and DepMap. Among 36 protein prediction models built by EN, the mean of Spearman rank correlation is 0.360 (95% CI: [0.281-0.439]) for external validation, and the prediction of 75% of the proteins is significantly correlated with the real expression (p value < 0.05) (Supplementary Table). In general, a model with a better performance in the training set leads to a better prediction in the external set (correlation of 0.48, p < 0.01). While excluding phosphorylated/cleaved proteins, the correlation could reach 0.42. Interestingly proteins including alpha-Catenin, N-Cadherin, beta-Catenin, p53_Caution could achieve remarkable performance (Spearman > 0.7). We further inspected the top poorly predicted proteins ER-alpha and PR and found the correlation of their actual expression between DepMap and CellMinerCDB is only 0.035 and 0.25, respectively. The undesired performance reinforces the importance of high-quality data for both training and testing. In addition, the slight decrease of predictive performance is likely due to the batch effect of gene expression profiles and the lack of optimal normalization of the external data in the model. Nevertheless, under these various challenges, the independent external validation for the models of those shared proteins still provides reasonable results.

Next, we turned to the recent DepMap 20Q1 release to explore if the model is applicable to prospective data. We found 36 new cell lines with both gene expression and gene effect scores. For each cell line, we used TransCell to predict the effect score of 2000 genes. The predicted effect scores are significantly correlated with the real effect score, with over 70% of cell lines presenting a high Spearman correlation coefficient of > 0.7 (Figure 7a). Besides, we investigated three new pediatric cancer cell lines. As expected, all of them have a correlation higher than 0.7 (Figure 7b). Given the finding that the correlation of all the gene scores across all genes and cell lines between two independent experimental CRISPR-Cas9 screen studies could only achieve 0.7 [38], the remarkable performance of TransCell warrants its wide application in other cell lines.

**Figure 7.**
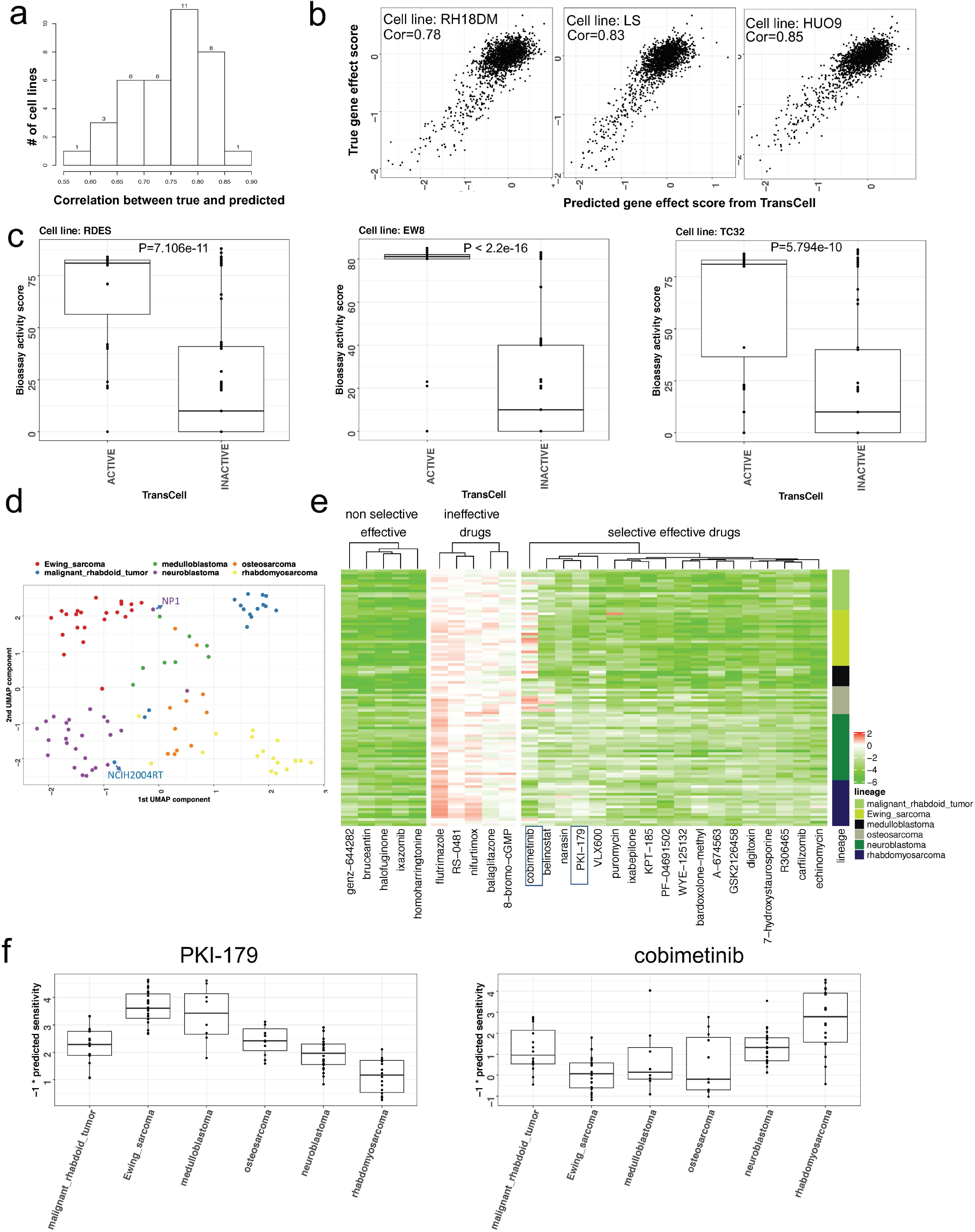
*In silico* expanding measurements for pediatric cell lines. **a**. Distribution of correlations between predicted and true gene effect scores across 36 new cell lines. **b**. Scattering plots of predicted gene effect scores and true gene effect scores in three new pediatric cell lines. **c**. Validation of TransCell drug sensitivity prediction using published screening results in three pediatric cell lines. Active was defined as TransCell score < -2. The active score provided in PubChem Bioassay was adopted for comparison, and the two-tailed student t-test was used to compute the difference. **d**. UMAP visualization of 101 pediatric cell lines based on predicted sensitivity of 2,356 drugs. Only the cancer with at least ten cell line profiles was selected. **e**. Heatmaps of predicted sensitivity of non-selective effective drugs, ineffective drugs, and selective effective drugs. Non-selective drugs: highly effective in the majority of cell lines (score < -2); ineffective drugs: ineffective in the majority of cell lines (score > -2); selective-effective drugs: only effective in a specific cancer. For each cancer, a fisher-exact test was used to compute compound selectivity. Only five non-selective drugs and five ineffective drugs were visualized. **f**. Example of two selective drugs PK1-179 and cobimetinib.

#### Case 1: *In silico* drug screening for pediatric cancer cell lines identifies new repurposing candidates

We implemented TransCell to infer the sensitivity of 4,686 DepMap drugs for the 124 pediatric cell lines recently published [33] (Supplementary Table). We started with evaluating the performance in three Ewing Sarcoma cell lines (RDES, EW8, TC32), for which we could find the screening results in PubChem Bioassay. The sensitivity data of 305 drugs in each cell line were retrieved from PubChem. As expected, the active compounds predicted by TransCell (score < -2) have much higher activity than inactive compounds in all cell lines (Figure 7c), further confirming its reasonable performance. Next, we clustered all the cell lines of six common pediatric cancers and found the predicted drug sensitivity could further classify cancer types (Figure 7d). Interestingly, we observed that some cell lines have very different responses from most of the cell lines with the same origin. For example, NB1 (neuroblastoma cell line) and NCIH2004RT (malignant rhabdoid tumor) more closely resemble Ewing Sarcoma and neuroblastoma, respectively. We further investigated individual drugs. In total, 29 drugs including homoharringtonine and ixazomib are effective in over 90% of pediatric cancer cell lines (Figure 7e). Notably, a number of drugs are only effective in specific cancers: neuroblastoma (8 drugs), Ewing sarcoma (7), rhabdomyosarcoma (3), medulloblastoma (1), malignant rhabdoid tumor (0), and osteosarcoma (0) (Figure 7e). For instance, Ewing sarcoma and medulloblastoma cells are more sensitive to PKI-179, a PI3K/mTOR inhibitor, while rhabdomyosarcoma cells are more sensitive to cobimetinib, a MEK inhibitor (Figure 7f). The prevalence of selective compounds implicates the potential of leveraging the predicted profiles to identify novel repurposing candidates and their biomarkers.

#### Case 2: *In silico* CRISPR knockout captures drug resistance in melanoma

Existing cell lines are frequently modified such as drug treatment in resistance mechanism elucidation and gene knock-out in therapeutic target identification, leading to many variations of cell lines that may present different molecular profiles from the parent cell lines. A survey of the RNASeq repository ARCHS4 [39] indicated that some cell lines even have thousands of gene expression profiles (Figure 8a). Obtaining comprehensive molecular measurements would greatly assist the research; however, the high cost of experiments does not allow to do so most of the time. Here, using melanoma-resistant cell lines as an example, we demonstrate TransCell could provide such data efficiently. In this study [40], to understand dynamic transcriptomic states, the authors profiled parental cell lines (including M229 and M238) treated with DMSO/vehicle, on-treatment lines (days to weeks on BRAFi), and resistant lines (months to years on BRAFi or BRAFi+MEKi). Using their gene expression profiles (access from Gene Expression Omnibus (GEO) GSE75299), we deployed TransCell to predict the BRAF gene effect score. It is known that the cells that were treated with BRAFi became less dependent on BRAF. TransCell could nicely capture the difference of BRAF dependencies among three groups (pretreatment, on-treatment, resistant) in both M229 and M238 (Figure 8b). The predicted scores of our non-resistant lines are close to the scores in the majority of skin cancer cell lines [4] and the predicted scores of the resistant M229 line are similar to the scores of another resistant M229 line for which our lab profiled recently for drug discovery [41] (The predicted scores of the first three resistant samples in GEO GSE145990 are -0.58, -0.54, -0.52). Although these experiments were conducted in different labs, the predicted scores were very consistent. Together, we show that TransCell gene effect prediction could be a proximity of the CRISPR-Cas9 knock-out experiment when a genome-wide knockout experiment is not feasible to perform.

**Figure 8.**
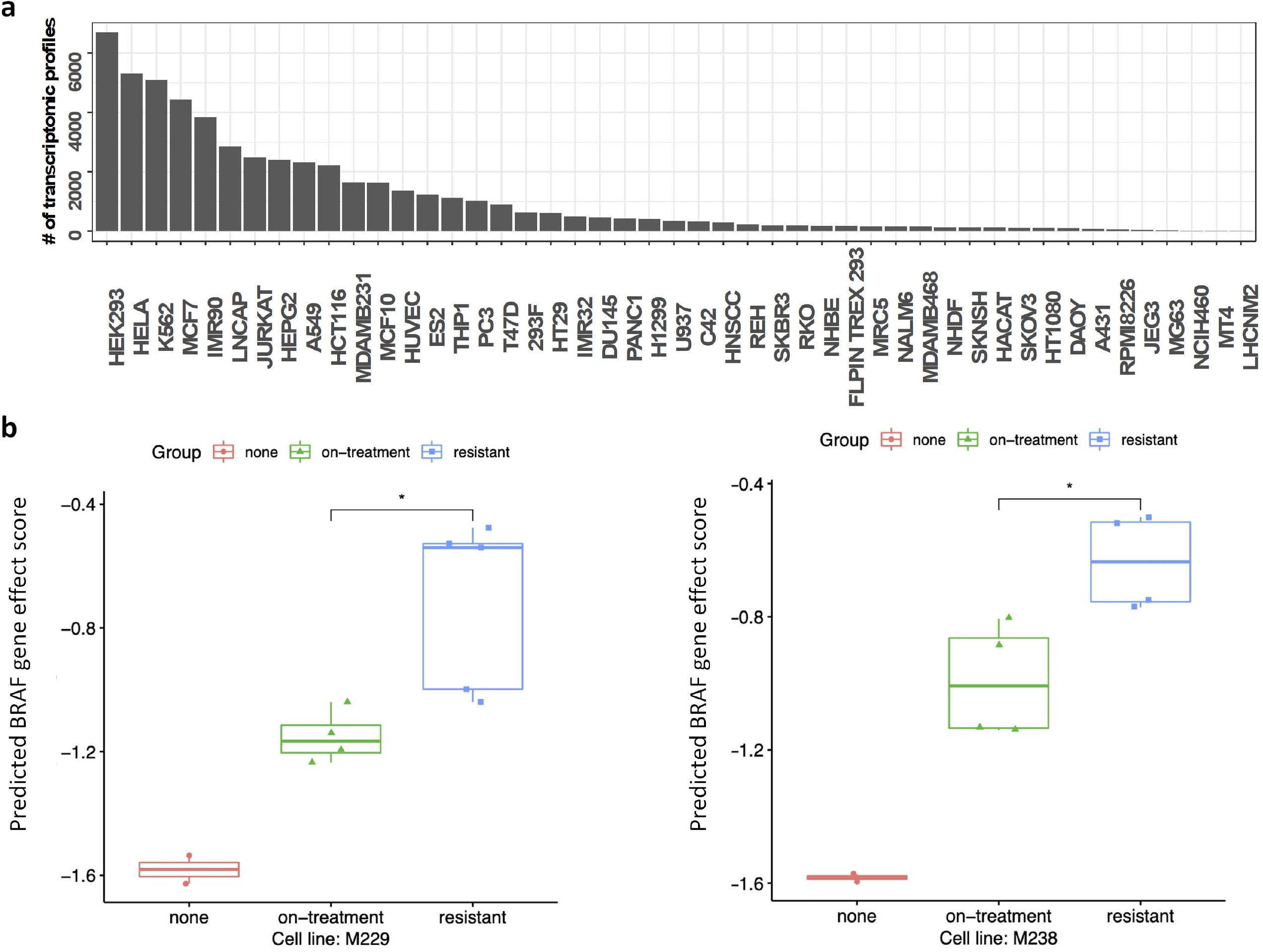
*In silico* CRISPR knockout captures drug resistances. **a**. The number of RNASeq profiles available per cell line. The numbers were collected from the ARCHS4 website (https://maayanlab.cloud/archs4/). **b**. Predicted BRAF gene effect score in two melanoma cell lines M229 and M238. RNASeq profiles downloaded through GEO (GSE75299) were processed using the OCTAD pipeline (Methods). A lower score means a high gene dependency (DepMap used -1 as a threshold to define the dependency). A two-sided student t-test was used to compute the difference. Note that the top sample in the resistant group in M229 was treated by the combination of both BRAF and MEK inhibitors.

### TransCell: a web portal to predict six measurement types solely based on gene expression profiles

To have a broad impact to the community, we develop TransCell into a R-Shiny web portal that allows researchers predicting six measurement types including metabolite, protein, CNV, gene effect score, drug sensitivity, and mutation given gene expression profiles. It is noted that we trained one model for each individual feature and searched optimal hyperparameters for each model through Keras Tuner. Furthermore, for each individual feature, in order to obtain robust results, we saved five models based on 5-fold cross validation. TransCell later would utilize them to provide the average predicted result. In total, we have saved 352,740 models ([225 (# of metabolites) + 214 (# of proteins) + 18,333 (# of genes in gene effect score dataset) + 4,686 (# of drugs in drug sensitivity dataset) + 27,639 (# of genes in CNV dataset) + 19,451 (# of genes in mutation dataset)]*5 = 352,740). They take around 4 terabytes on the Amazon AWS server. The training time of 5-fold cross validation for each individual feature is about five minutes. All the models were processed by GNU parallel computation [42]. The interface of TransCell web portal is shown in Figure 9. Users could upload new gene expression profiles (log2(1+tpm)) and select their interested features for making predictions. After TransCell computation is finished, users can either download the results from the portal or can get it through email if valid email address is provided.

**Figure 9.**
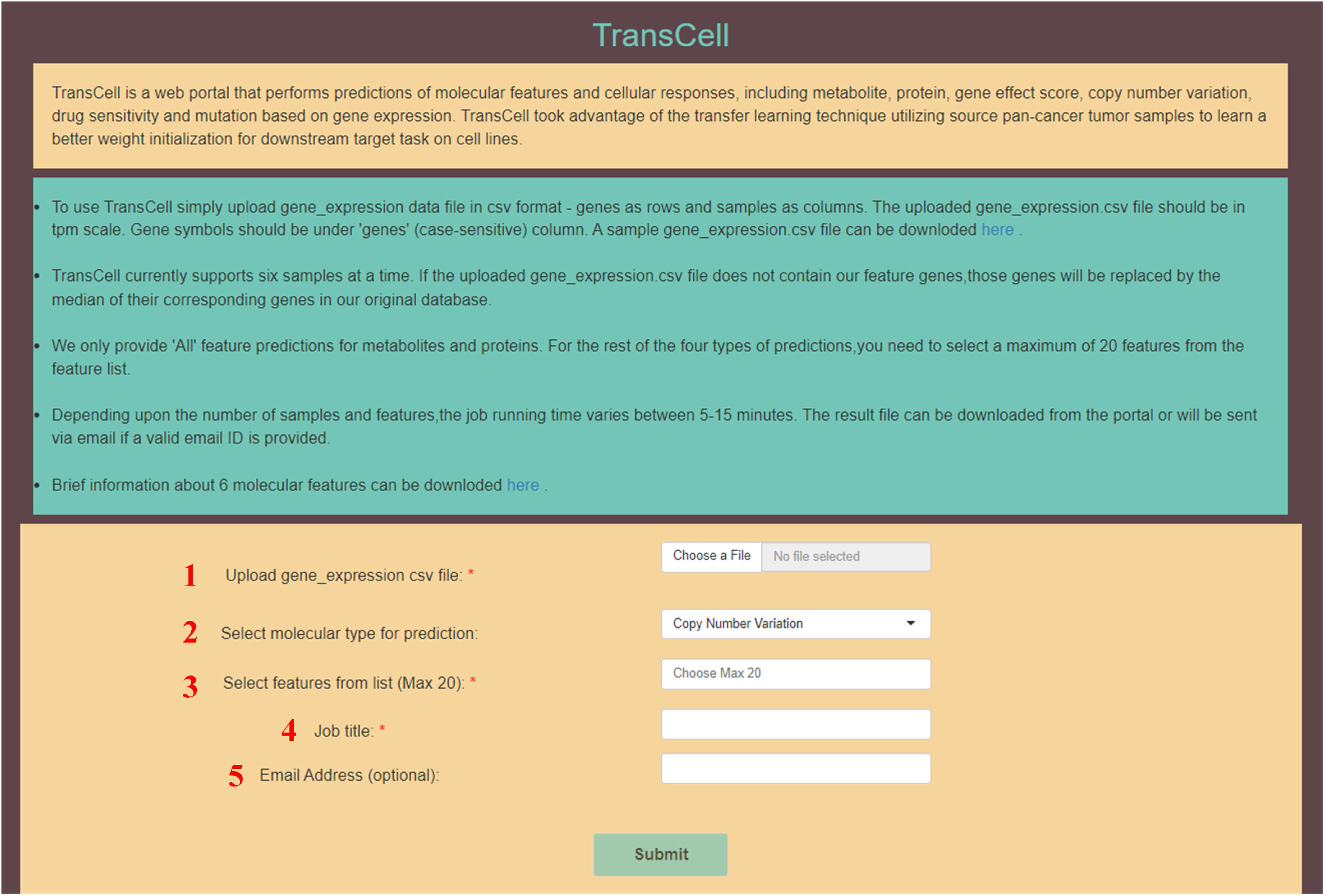
TransCell web portal. 1: Upload users’ gene expression CSV profile. 2: Select the measurement type for making prediction. 3: Select features from drop down menu. 4: Give a job title. 5: Provide an email address (optional).

## DISCUSSION

Gene expression is arguably the most widely used modality in biomedical research. GEO, the largest expression repository, has archived 42 million profiles as of Jan 2021, and the number continues to multiply. The advances of single-cell technology even enable the generation of thousands of expression profiles in a single experiment. Gene expression profiling of new samples today becomes routine. Compared to gene expression, other modalities, especially those large-scale perturbation experiments, are not easy, if not impossible, to produce in most labs. If gene expression could be leveraged to predict other modalities, additional lenses could be added to gain more biological insights. It is known that gene expression encodes the information of other modalities, but whether the baseline models and existing data could decode such relations remains unknown. To our best knowledge, this work is one of the first to provide a comprehensive study of molecular measurement prediction using gene expression data. We also developed TransCell, which utilizes the transfer learning technique to gain better weight initializations learned from a larger dataset TCGA, for improving the performance of a model.

Except for genes with data imbalanced issues in mutation prediction, the easiness of predicting six measurement types is ranked as follows: mutation, CNV, protein, metabolite, gene effect score, and drug sensitivity. The top performance of mutation, CNV, and protein levels could reflect their inherent relationship that genetic variation regulates gene expression, which controls protein abundance. Metabolite abundance is directly related to protein expression rather than mRNA expression, and its abundance is very dynamic; however, surprisingly, the majority of metabolites can be fairly predicted using TransCell. The inferior performance of perturbation experiments (CRISPR knockout and drug treatments) confirms the complicated cellular responses that could not be easily captured by linear models. Besides, the quality of the high throughput experiments sets back model development [43]. TransCell adequately addresses such challenges. Notably, TransCell performs reasonably well in prospective datasets as well as external datasets. Two case studies further suggest the potential of using TransCell in biomedical research. Anticipating the fast growing perturbation experiments in vitro cell lines (e.g., drug screening, CRISPR genome-wide knockout), we develop the TransCell web portal as a generally applicable prediction framework.

As protein is the direct product of mRNA, the remarkable predictive power is not surprising. The correlation between gene expression levels and protein levels is varying with respect to cell-state transitions. The steady-state represents the cells undergoing long-term dynamic processes, such as continuous proliferation, differentiation, or other types of fate decisions [44, 45]. During the steady state, the mRNA levels primarily determine protein abundance. Their linear relation may explain the considerable performance of linear-based models. The least predictable proteins in the scale-up protein prediction tend to belong to a protein complex. This could be explained by the fact that the degradation rates of proteins can be different if they are in a complex [46]. Differences in degradation rate may decrease the correlation between mRNAs and proteins. The gene enrichment analysis of biological processes for the poorly predicted proteins uncovered most enriched pathways are related to the response of stimulus. The short-term adaptation of cells to new states or environmental conditions change often comes with post-transcriptional mechanisms. The short-term adaptation makes the delay between transcriptional induction and protein level increase [47]. In short, the low mRNA-protein correlation results in poor predictions for those proteins under the aforementioned scenario.

Genetic structural variation in the human genome can be presented in many forms, ranging from single nucleotide polymorphisms to CNV [48]. Changes in copy number might alternate gene expression levels [49]. Gene-wise copy number amplification tends to enhance gene expression, while copy number deletion usually leads to decreased gene expression. Taking linear regression into account in analyzing the relationship between CNV and expression level, plenty of genes had a high degree of fitting, confirming that gene expression change directly reflects CNV in many cases [50].

In mutation prediction, we only labeled two classes denoting whether a cell line harbors a given mutation or not. The top 20 selected genes have at least 21.7% of cell lines harboring a mutation. To the scale-up mutation prediction, the top 2000 selected genes at least have 5% of cell lines with a mutation. TransCell does not perform as well as Logistic_LASSO, but the performance is reasonable (with ∼0.9 of AUC). We did not take the classic imbalance problem into account in order to make a consistent comparison across all the measurements. For genes with lower mutation proportion across all cell lines, the model would have poor predictive performance, especially for the minority class. In TransCell web portal, we only provide 15010 models which macro-f1 scores are greater than 0.7 in mutation prediction. Nevertheless, this survey confirms the potential power of using gene expression to infer mutations.

The high dimensional gene expression features and limited cell lines pose challenges to any machine learning algorithms [51]. Prior studies show that autoencoder, an unsupervised artificial neural network that could capture non-linear relationships between features, can learn a robust representation of gene expression data. We also observed the superiority of autoencoder in representing tumor gene expression profiles in reference tissue selection [52]. However, its performance highly depends on the initial parameters when the training set is small. One approach to overcome data scarcity is to apply transfer learning techniques. Transfer learning aims to improve the performance of target learners on target domains by transferring the knowledge learned from different but related source domains and then apply the knowledge to a target task [53, 54]. The weights and architecture obtained from the pre-trained model could be directly used to initialize the target task [55]. Moreover, weight optimization in non-linear autoencoders is cumbersome. With large initial weights, autoencoders typically find poor local minima. A pre-training procedure was introduced to provide an effective way of initializing the weights of deep autoencoder networks [56]. Here, TransCell confirms that transfer learning technique can improve model prediction performance through pre-training.

We found that the two-step pre-trained CCLE_enc_ outperformed the other two types of encoders. TransCell took advantage of the transfer learning scheme learned on a different but related system to compensate for the small dataset issue and improved the model performance by obtaining better weight initializations. Although TCGA does not have metabolite and perturbation data, since it aids gene expression feature representation, the resulting features have superior performance regardless of molecular types. Moreover, without the feature embedding step using an autoencoder, the simple combination of PCA and DNN performs poorly across all the comparisons, suggesting the importance of feature selection in this task.

Although we have demonstrated the feasibility of using gene expression to predict other measurements, the following work could be conducted in the future. First, the meta data of cell lines and tumors including age, gender, and disease lineage could contribute to model performance. We conducted experiments in which commonly used meta data including age, gender, and disease lineage were added as features in the metabolite prediction; we did not observe any significant improvement (Supplementary Text and Figures S4-S5). Even though some meta data might be informative in some tasks, the scarcity of meta data associated with gene expression profiles limits its power in practice. Second, the evaluation of TransCell using external profiles is highly desired in order to confirm the robustness of this approach. In particular, due to the differences between cell lines and tissues [57, 58], additional care should be taken while applied to those samples different from cancer cell lines. Interestingly, in additional exploration of using the TransCell concept to predict clinical responses, we observed that a two-step deep transfer learning model performed better than that without transfer learning (Supplementary Text and Figure S6). Third, the performance of TransCell in the prediction of complicated measurements such as metabolite could be improved through further tuning and integration with more biological data such as protein-metabolite and protein-protein interactions. We used Keras tuner to search for the optimal hyperparameters, while the interpretation of individual performances in poorly predicted measurements and a large-scale comparison of various hyperparameters in individual tasks could help improve the models. Moreover, a multi-task learning model for metabolite predictions based on TransCell’s architecture was built; however, sharing information with unrelated task might result in negative transfer (Supplementary Text and Figure S8-S9). Given the nature of the metabolic profiling methods, the abundance of different metabolites cannot be compared [26], thus additional care is needed while attempting to leverage their relations in modeling. Lastly, different from many published studies where additional omics data other than gene expression such as chemical structures and mutations were incorporated in order to improve performance [59]. This work aims to answer if it is feasible to use gene expression alone to predict other measurement types; meanwhile, a user-friendly service, TransCell web portal is available where six molecular measurement types can be predicted from a single website. In the potential applications, acquiring additional data for new gene expression profiles is challenging, if not impossible, thus our model comparison tended not to include other additional features, increasing the difficulty of comparing with the state-of-the-art models that often need additional features. We indeed compared one relevant published model DeepDR [9] in drug sensitivity prediction and demonstrated the superiority of TransCell (Supplementary Text and Figure S7). In short, this survey confirms that large-scale *in silico* characterization of genomic landscape and cellular responses from gene expressions for new cell lines is feasible.

## Supporting information

Supplementary Doc

## Code availability

All the codes required in this study are available in GitHub (https://github.com/Bin-Chen-Lab/transcell). The models were evaluated under AWS g3.4xlarge.

## Author contributions

B.C. conceived and supervised the study; S.Y. developed the method and performed the analysis with the input from J.X., M.S., K.L., S.P., R.C., J.Z. and B.C.; R.C. led external validation. S.P. developed the web portal. S.Y. and B.C. wrote the manuscript with input from J.X., M.S., and J.Z..

## Acknowledgments

We would like to thank Chen lab members for their critical comments. The research is supported by R01GM134307, K01 ES028047, and the MSU Global Impact Initiative. The content is solely the responsibility of the authors and does not necessarily represent the official views of sponsors.

