## Supplementary Doc for "TransCell: In silico characterization of genomic landscape and cellular responses from gene expressions through a two-step deep transfer learning"

### Supplementary Materials

#### The architecture of TransCell for metabolite prediction

Here, we provide the detailed architecture of TransCell in metabolite prediction. P used Adam as an optimizer (learning rate = 0.001) with MSE loss. The input layer merging the output of two-step pre-trained CCLEenc had 352 neurons, followed by two hidden layers with 320 and 64 neurons. The activation function was set as a hyperbolic tangent for each layer. Moreover, to mitigate the overfitting problem, except for the output layer, the regularization penalties were applied to the output and bias for each layer. We used dropout 0.55, 0.60, and 0.50 on the input layer and two hidden layers, respectively. In addition, early stopping was used to terminate model training once the model performance stops improving on the validation set. The optimal hyperparameters in TransCell were searched by the Keras Tuner with 100 max trials. Before computing the evaluation metrics and Spearman rank correlation, we performed the inverse transformation on the predicted values, making them back to their original space without data rescaling by min-max scaler.

#### The architecture of TransCell for gene effect score prediction

In gene effect score prediction, the cancer cell line RNA-seq TPM gene expression data (CCLE) was rescaled in the range of 0 to 1 and the target data was rescaled in the range of -1 to 1 by min-max scaler. The TransCell contains two-step pre-trained CCLE encoder ($CCLE_{enc}$) and a feed forward prediction network (P). For the architecture of P, we used Adam as an optimizer (learning rate = 0.0009) with mean squared error loss. The input layer merging the output of two-step pre-trained $CCLE_{enc}$ had 288 neurons, followed by two hidden layers with 128 and 64 neurons. The activation function was set as a hyperbolic tangent for each layer. Moreover, to alleviate model suffering from overfitting, except for the output layer, we added L2 regularization penalties on the output and bias for each layer. The dropout was 0.60, 0.35, and 0.2 for the input layer and two hidden layers, respectively. In addition, early stopping was used to terminate model training once the mode performance stops improving on the validation set. The optimal hyperparameters in the TransCell were searched by Keras Tuner with 100 max trails. Before computing the evaluation metrics and Spearman rank correlation, we performed an inverse transform on the predicted values.

#### The architecture of TransCell for drug sensitivity prediction

In drug sensitivity prediction, the CCLE was rescaled in the range of 0 to 1 and the target data was rescaled in the range of -1 to 1 by min-max scaler. The TransCell is comprised of two-step pre-trained $CCLE_{enc}$ and P. For the architecture of P, we used Adam as an optimizer (learning rate = 0.0005) with mean squared error loss. The input layer merging the output of two-step pre-trained $CCLE_{enc}$ had 256 neurons, followed by two hidden layers with 192 and 96 neurons. The activation function was set as a hyperbolic tangent for each layer. Moreover, to address the overfitting problem, except for the output layer, the L2 regularization penalties were applied to the output and bias for each layer. We added dropout 0.50, 0.30, and 0.20 for the input layer and two hidden layers, respectively. In addition, early stopping was implemented in the training process to stop model training once the loss of validation starts to increase. The optimal hyperparameters in the TransCell were searched by Keras Tuner with 100 max trails. Before computing the evaluation metrics and Spearman rank correlation, we performed inverse transform on the predicted values.

#### The architecture of TransCell for protein prediction

In protein prediction, the CCLE was rescaled in the range of 0 to 1 and the target data was rescaled in the range of -2 to 2 by minmax scaler. The TransCell is composed of two-step pre-trained $CCLE_{enc}$ and P. For the architecture of P, we used Adam as an optimizer (learning rate = 0.0001) with mean squared error loss. The input layer merging the output of two-step pre-trained $CCLE_{enc}$ had 512 neurons, followed by two hidden layers with 224 and 96 neurons. The activation function was set as a hyperbolic tangent for each layer. Moreover, to mitigate the overfitting problem, except for the output layer, the L2 regularization penalties were applied to the output and bias for each layer. The dropout was 0.55, 0.25, and 0.4 for the input layer and two hidden layers, respectively. In addition, we used early stopping in the training process. The optimal hyperparameters in the TransCell were searched by Keras Tuner with 100 max trails. Before computing the evaluation metrics and Spearman rank correlation, we performed an inverse transform on the predicted values.

#### The architecture of TransCell for copy number variation prediction

In copy number variation prediction, the CCLE was rescaled in the range of 0 to 1 and the target data was rescaled in the range of -1 to 1 by minmax scaler. The TransCell is made up of two-step pre-trained $CCLE_{enc}$ and P. For the architecture of P, we used Adam as an optimizer (learning rate = 0.001) with mean squared error loss. The input layer merging the output of two-step pre-trained $CCLE_{enc}$ had 256 neurons, followed by two hidden layers with 256 and 32 neurons. The activation function was set as a hyperbolic tangent for each layer. Moreover, to address the overfitting problem, except for the output layer, we added the L2 regularization on the output and bias for each layer. The dropout was 0.60, 0.35, 0.55 for the input layer and two hidden layers, respectively. Besides, we used early stopping to terminate the training process once the validation loss showing an increased trend. The optimal hyperparameters in the TransCell were searched by Keras Tuner with 100 max trails. Before computing the evaluation metrics and Spearman rank correlation, we performed an inverse transform on the predicted values.

#### The architecture of TransCell for mutation prediction

In mutation prediction, the CCLE was rescaled in the range of 0 to 1 by the min-max scaler. The TransCell consists of two-step pre-trained $CCLE_{enc}$ and P. For the architecture of P, we used Adam as an optimizer (learning rate = 0.001) with binary cross-entropy loss. The input layer merging the output of two-step pre-trained $CCLE_{enc}$ had 416 neurons, followed by two hidden layers with 192 and 32 neurons. The activation function was set as relu except for the output layer using the sigmoid function. Moreover, considering the overfitting problem, except for the output layer, we added the L2 regularization on weights for each layer. The dropout was 0.05, 0.30, 0.05 for the input layer and two hidden layers, respectively. In addition, we used early stopping in the training process. The optimal hyperparameters in the TransCell were searched by Keras Tuner with 100 max trails.

#### Feature Selection Methods

Going from microarray to next generation sequence has given rise to a wealth of feature selection and dimension reduction techniques [1]. Feature selection methods are generally divided into three categories: filter methods, wrapper methods, and embedded methods [2]. The filter methods such as mutual information, Pearson correlation criteria and Chi-square test are independent of any following learning algorithms. Due to the reduced computational time, filter methods are effective for high dimensional datasets; however, the nature that they consider each feature separately might weaken model performance [3]. Wrapper methods rely on learning algorithms to find a subset of features whose interactions would be considered during searching. Compared to filter methods, wrapper methods have higher computational costs and risk of overfitting [4, 5]. Embedded methods combine the advantages of both filter and wrapper methods. Selecting the feature subset is considered as a part of model construction, implemented in many learning algorithms such as Adaboost [6], random forest, and decision tree [7, 8]. The most common type of embedded methods is regularization methods, including least absolute shrinkage and selection operator (LASSO) regression [9], ridge regression [10], elastic net (EN) [11], $l_1$ regularized logistic regression [12], and logistic regression relying on the elastic net penalty [13].

Lasso regression performs $l_1$ regularization. The optimization problem is equivalent to the parameter estimation that follows:

$$\hat{\beta}(\lambda) = \underset{\beta}{\operatorname{argmin}}\left(\frac{\|Y - X\beta\|_2^2}{2n} + \lambda\|\beta\|_1\right) \tag{S1}$$

where $n$ means the number of samples, $\|Y - X\beta\|_2^2 = \sum_{i=0}^{n}(\mathrm{Y}_i - (\mathrm{X}\beta)_i)^2$, $\|\beta\|_1 = \sum_{j=1}^{k}|\beta_j|$, and $\beta_j$ is the coefficient for feature $j$. $\lambda \geq 0$ penalizes the coefficients of the regression variables and shrinks some of them to zero. In this way, the variables that still have a non-zero coefficient after the shrinking process are selected to be essential features. The EN adds an additional $l_2$ regularization term into LASSO loss function. The EN loss function is shown as below:

$$\hat{\beta}(\lambda) = \underset{\beta}{\operatorname{argmin}}\left(\frac{\|Y - X\beta\|_2^2}{2n} + \lambda_1\|\beta\|_1 + \lambda_2\|\beta\|_2^2\right) \tag{S2}$$

where $\lambda_1 = \alpha * l_{1-ratio}$ and $\lambda_2 = \frac{\alpha * (1 - l_{1-ratio})}{2}$, which $\alpha$ is a constant that multiplies the penalty term and $l_{1-ratio}$ is the elastic net mixing parameters with $l_{1-ratio} \in [0,1]$ deciding a combination of $l_1$ and $l_2$ penalty. For instance, if $l_{1-ratio} = 0$, the penalty is a $l_2$ regularization. If $l_{1-ratio} = 1$, the penalty would be $l_1$ regularization. In other words, the EN can be seen as a linear combination of the LASSO and Ridge penalty. Therefore, EN has similar behavior, shrinking model weights, as LASSO allows essential groups of correlated features to be selected. RF also offers a feature selection indicator. It uses variance of out-of-bag errors from permutations to compute the importance of each feature during the training process. Moreover, PCA is a dimensionality reduction technique that projects the data into a lower dimensional space and retains most of the variation part in the data set. By doing so, the most important information from the original data could be represented by fewer components. Moreover, in binary mutation data, Logistic_LASSO and Logistic_EN are applied for feature selection keeping informative features out of the original data by removing irrelevant and redundant ones. Given a dataset $X$ ($X = \{\mathrm{x}_1, ..., \mathrm{x}_n\}$), which has $n$

samples and *p* features, let $y \in \{0,1\}$ standing for the binary target of each sample. The conditional probability distribution of the class label *y*, given a feature vector *x*, is as below:

$$\text{Prob}(y \mid x) = \frac{1}{1+\exp(-y(w^T x + c))} \tag{S3}$$

where $x \in \Re^p$ denotes an observation made of *p* features. The weight vector $w \in \Re^p$ and intercept $c \in \Re$ are the parameters of the logistic regression model. The log-likelihood function associated with the learning samples $\{(x_i, y_i),\ i=1, ..., n\}$ based on the equation (S3) is defined as:

$$l(\text{w},\text{c}) = \sum_{i=1}^{n} \log(\exp(-y_i(w^T x_i + c)) + 1) \tag{S4}$$

A maximum likelihood estimation of the model parameters *w* and *c* would be obtained by minimizing the equation (S4). Therefore, for Logistic_LASSO, which employs a $l_1$ regularization term on the regression parameters, the loss function of the optimization problem is shown as below:

$$\min_{w,c} l(w,c) + \|w\|_1 = \min_{w,c} \sum_{i=1}^{n} \log(\exp(-y_i(w^T x_i + c)) + 1) + \|w\|_1 \tag{S5}$$

Moreover, to Logistic_EN, the logistic regression with a combination of $l_1$ and $l_2$ regularization terms, its loss function is given as follows:

$$\min_{w,\text{c}} l(\text{w},\text{c}) + \rho\|w\|_1 + \frac{1-\rho}{2}\|w\|_2^2 = \min_{w,c} \sum_{i=1}^{n} \log(\exp(-\text{y}_i(\text{w}^T \text{x}_i + \text{c})) + 1) + \rho\|w\|_1 + \frac{1-\rho}{2}\|w\|_2^2 \tag{S6}$$

where $\rho \in [0,1]$ controls the strength of $l_1$ and $l_2$ regularizations.

#### Different feature set combination comparisons

Considering meta data for the clinical dataset (TCGA) and cell line dataset (CCLE), we run the TransCell to build all metabolite models under different feature set combinations (original features v.s. original features + age + gender) in the first experiment. The first one is the original feature set which we used to train the TransCell (5000 genes); and the second one is the feature set including 5000 genes, age, and gender features. To the gender feature, we give 1 to female and 0 to male. For the age feature, at first, we separate the age information into four groups (ex. 0-16, 16-45, 46-65, >65). Each group is transformed by one-hot encoding. By doing so, the feature size would be increased to 5005 features. From the experimental results in Figure S4a-c, we could find that two different feature sets lead to similar performance for TransCell metabolite predictions. The Wilcoxon rank sum test p-values are not significant for mean squared error, root mean squared error, and Spearman rank correlation.

Afterward, we take disease lineage, cell type information, into account when conducting the second experiment (original features v.s. original features + cell type features). There are 50 cell types from the union set in TCGA and CCLE datasets. We neglect samples with unknown cell type in CCLE. Therefore, there are 1429 and 15028 samples for CCLE and TCGA, respectively. The cell type information is encoded by one-hot encoding. We add additional 50 features in our original feature set. Therefore, in total, there are 5050 features. Compared to the original feature set which we used to train the TransCell (5000 genes), we find that adding cell type information would not change a lot toward the training performance for TransCell metabolite predictions as shown in Figure S5a-c. Meanwhile, the Wilcoxon rank sum test p-values are not significant for mean squared error, root mean squared error, and Spearman rank correlation as well. Overall, age, gender, and cell type information are not informative in this case.

#### Further application on clinical response prediction based on the TransCell concept

To evaluate the two-step transfer learning model on the clinical response prediction, we implemented a pre-two-stage transfer learning framework. We firstly learnt an encoder from the CCLE dataset, then transferred encoder's weights to the second autoencoder as its weight initialization. The second autoencoder was trained by the pan-cancer TCGA dataset. It is noted that the training features are 5000 genes, which are the same as TransCell. Afterward, we extracted the two-step pre-trained encoder to link to a prediction feedforward network (P). Our goal here is to use this framework, which is similar to TransCell, to predict clinical Cisplatin drug response based on the TCGA dataset. The Cisplatin TCGA dataset is from [14]. Based on the RECIST standard [15], we considered the clinical responses as two types, namely responder (including complete response and partial response) and non-responder (including stable disease and progressive disease). Therefore, it is a binary classification problem. Moreover, after removing the records of those patients who responded inconsistently to one drug during the course of treatment, there are 279 samples for Cisplatin drug response prediction.

Similar to the TransCell, this framework comprises a two-step pre-trained encoder and P. For the architecture of P, we used Adam as an optimizer (learning rate = 0.0005) with binary cross entropy loss. The input layer merging the output of two-step pre-trained encoder had 352 neurons, followed by two hidden layers with 224 and 64 neurons. The activation function was set as ReLU except for the output layer using the sigmoid function. Meanwhile, we added dropout 0.30 and 0.40 for the first and second layer of P, respectively. In addition, early stopping was implemented in the training process to stop model training once the loss of validation starts to increase. The optimal hyperparameters in this framework were searched by Keras Tuner with 100 max trails. In Figure S6, the AUCs are computed based on the five-fold cross validation. For the clinical Cisplatin drug response prediction, the framework built under TransCell's concept has the average of AUC 0.921. However, based on the same architecture, if we train the model directly without pre-two-stage transfer learning, the average of AUC would be 0.615, suggesting the importance of the transferred parameters.

#### Cancer cell line (CCLE) drug response prediction comparison between TransCell and DeepDR based on genomics of drug sensitivity in cancer (GDSC) project

Deep learning model to predict drug response (DeepDR) based on mutation and gene expression profiles of cancer cell or a tumor was proposed by Chiu et al. [16]. DeepDR model applied gene expression and mutation data to predict drug response (log-scale $IC_{50}$) for cancer cell lines (CCLE) based on Genomics of Drug Sensitivity in Cancer (GDSC) project [17]. Their model contains (i) a mutation pre-trained encoder trained by TCGA (ii) a gene expression pre-trained encoder trained by TCGA (iii) a drug response predictor network integrating the first two subnetworks. We refer to [16] for setting DeepDR architecture. To TransCell, it consists of a two-step pre-trained CCLE encoder and a prediction network (P) (please refer to Materials and Methods). TrasnCell was deployed to do the same task as DeepDR did in [16]. Therefore, we made drug response predictions for 265 drugs from the GDSC project by TransCell and DeepDR, respectively. Figure S7 suggests that TransCell has better performance. The RMSE of TransCell is significantly lower than DeepDR ($p = 5.2*10^{-6}$, Wilcoxon rank sum test).

#### Multi-task learning based on TransCell's architecture for metabolite predictions

Multi-task learning (MTL) aims at improving generalization by learning multiple tasks simultaneously. The MTL model transfers knowledge from one task to another wherever these tasks are related. It has been commonly utilized to address high dimensionality and small cohort size challenges. We further built a MTL model for metabolite predictions based on TransCell's architecture as shown in Figure S8. However, sharing information with unrelated task might hurt performance. Compared to the MTL model, TrasnCell has lower root mean squared error with significant p-value under the same 20 metabolite predictions as shown in Figure S9. In other words, without the prior knowledge of the relationship between metabolites, we observe negative transfer. Since the metabolites which users want to predict by TrasnCell web portal may not all be related. Therefore, we did not consider the MTL model in our case.

### Supplemental Figures

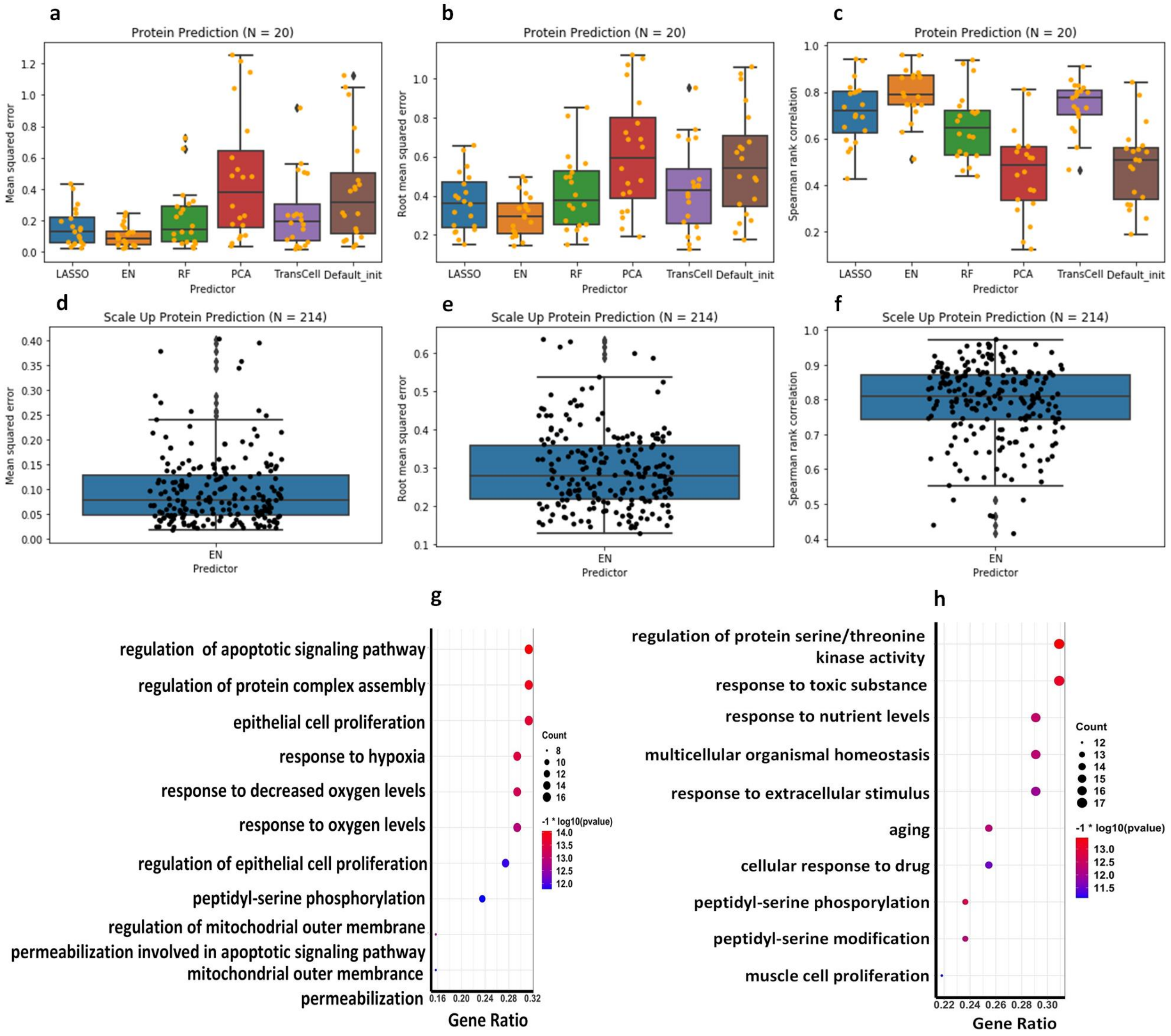


**Figure S1. Protein predictions. a-c.** The boxplots of (**a**) MSE, (**b**) RMSE, and (**c**) Spearman rank correlation for different predictors, including LASSO, EN, RF, TransCell, and two DNN designs with PCA and default initializations in 20 protein prediction models. **d-f.** The boxplots of (**d**) MSE, (**e**) RMSE, and (**f**) Spearman rank correlation for EN in all 214 scale-up protein prediction models. **g-h**. The dot plots of gene enrichment analysis of biological processes for target genes of proteins with RMSE (**g**) lower than the first quartile (well-predicted proteins) and (**h**) higher than the third quartile (poorly predicted proteins) in the scale-up protein predictions.

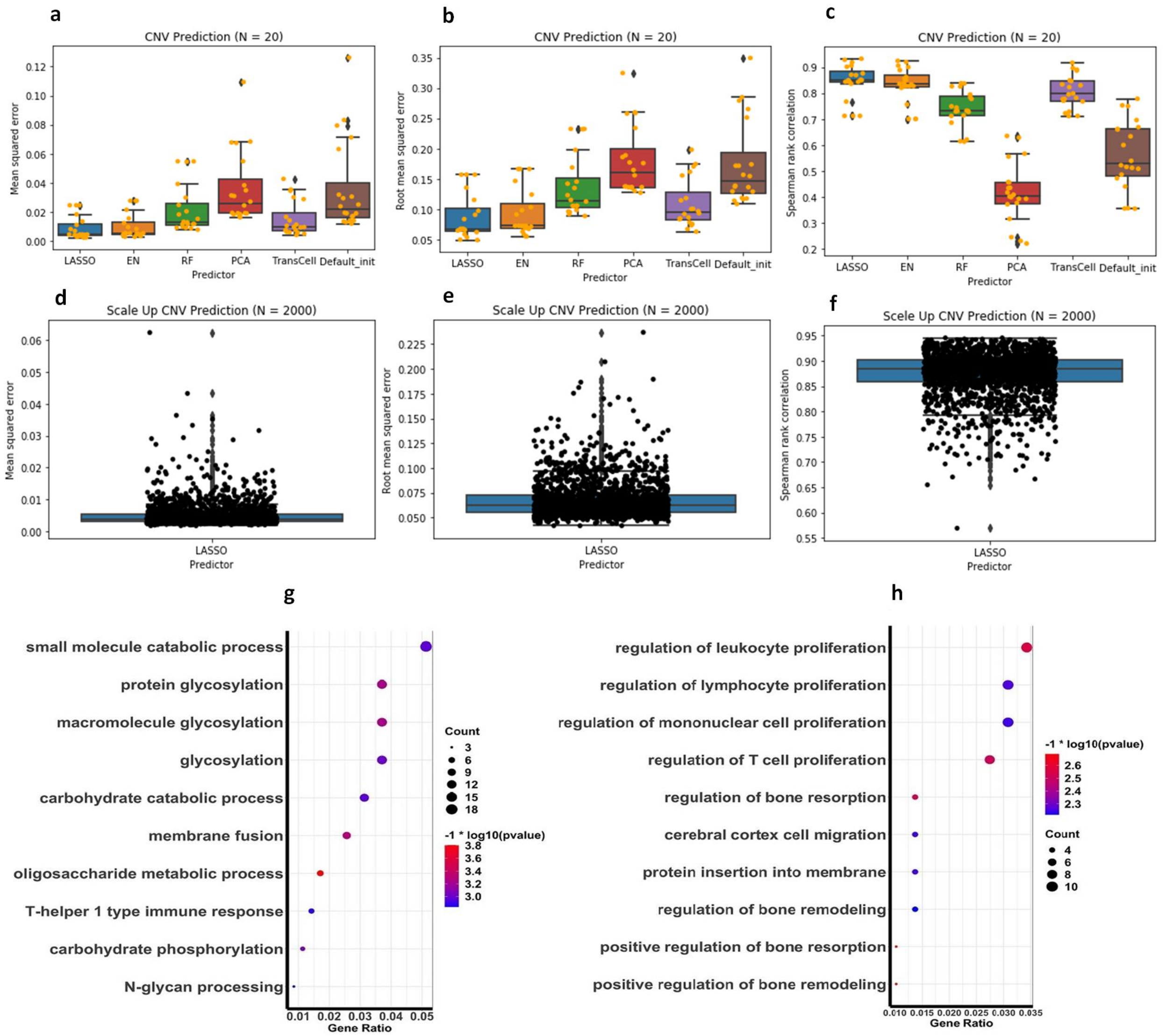


**Figure S2. CNV predictions. a-c.** The boxplots of (**a**) MSE, (**b**) RMSE, and (**c**) Spearman rank correlation for different predictors, including LASSO, EN, RF, TransCell, and two DNN designs with PCA and default initializations in 20 CNV prediction models. **d-f.** The boxplots of (**d**) MSE, (**e**) RMSE, and (**f**) Spearman rank correlation for LASSO in 2000 scale-up CNV prediction models. **g-h**. The dot plots of gene enrichment analysis of biological processes for genes with RMSE (**g**) lower than the first quartile (well- predicted genes) and (**h**) higher than the third quartile (poorly predicted genes) in the scale-up CNV predictions.

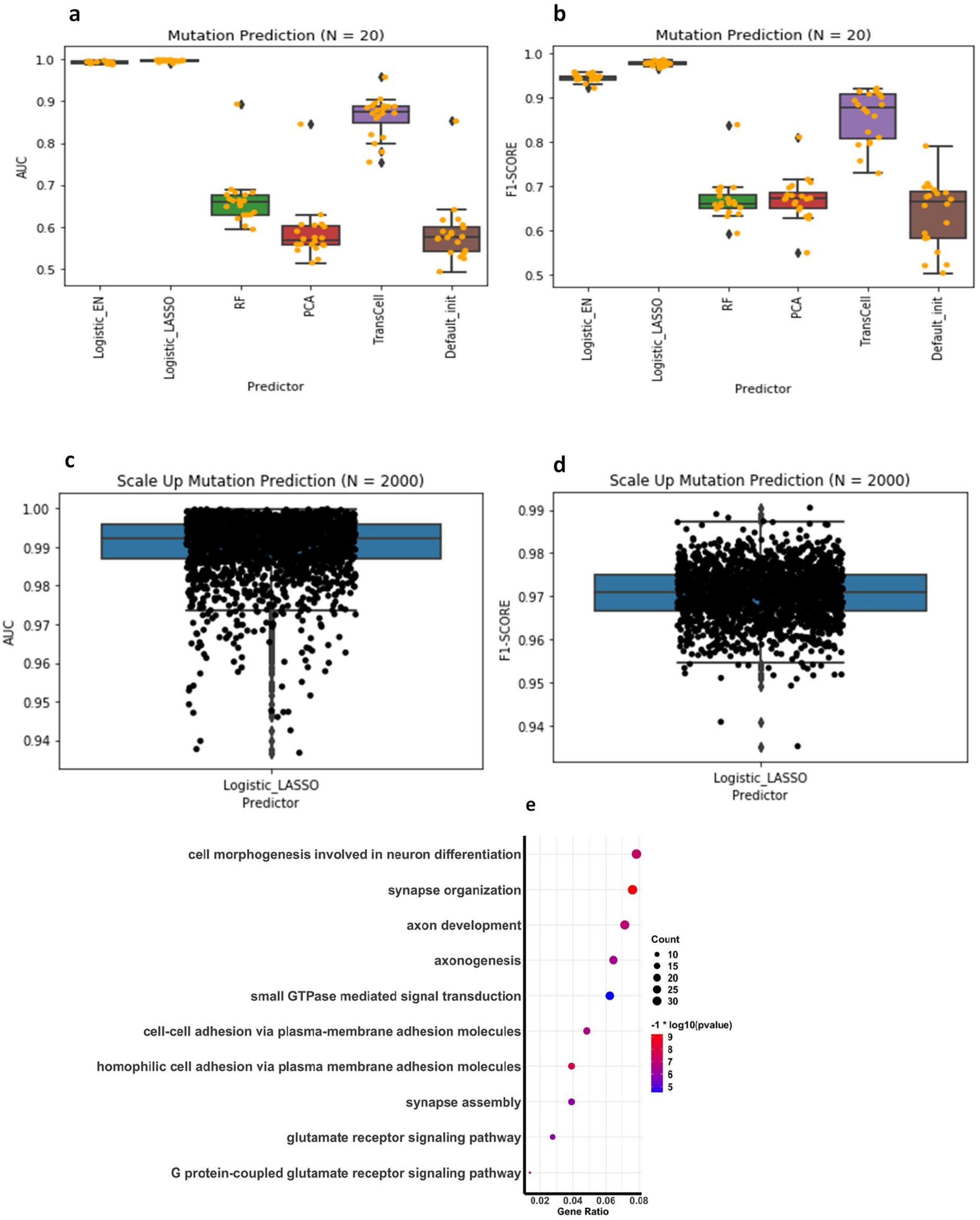


**Figure S3. Mutation predictions. a-b.** The boxplots of (**a**) AUC and (**b**) F-1 score for different predictors, including Logistic_EN, Logistic_LASSO, RF, TransCell, and two DNN designs with PCA and default initializations in 20 mutation prediction models. **c-d.** The boxplot of (**c**) AUC and (**d**) F-1 score for Logistic_LASSO in 2000 scale-up mutation prediction models. **e**. The dot plot of gene enrichment analysis of biological processes for genes with AUC lower than the first quartile (poorly predicted genes) in the scale-up mutation prediction.

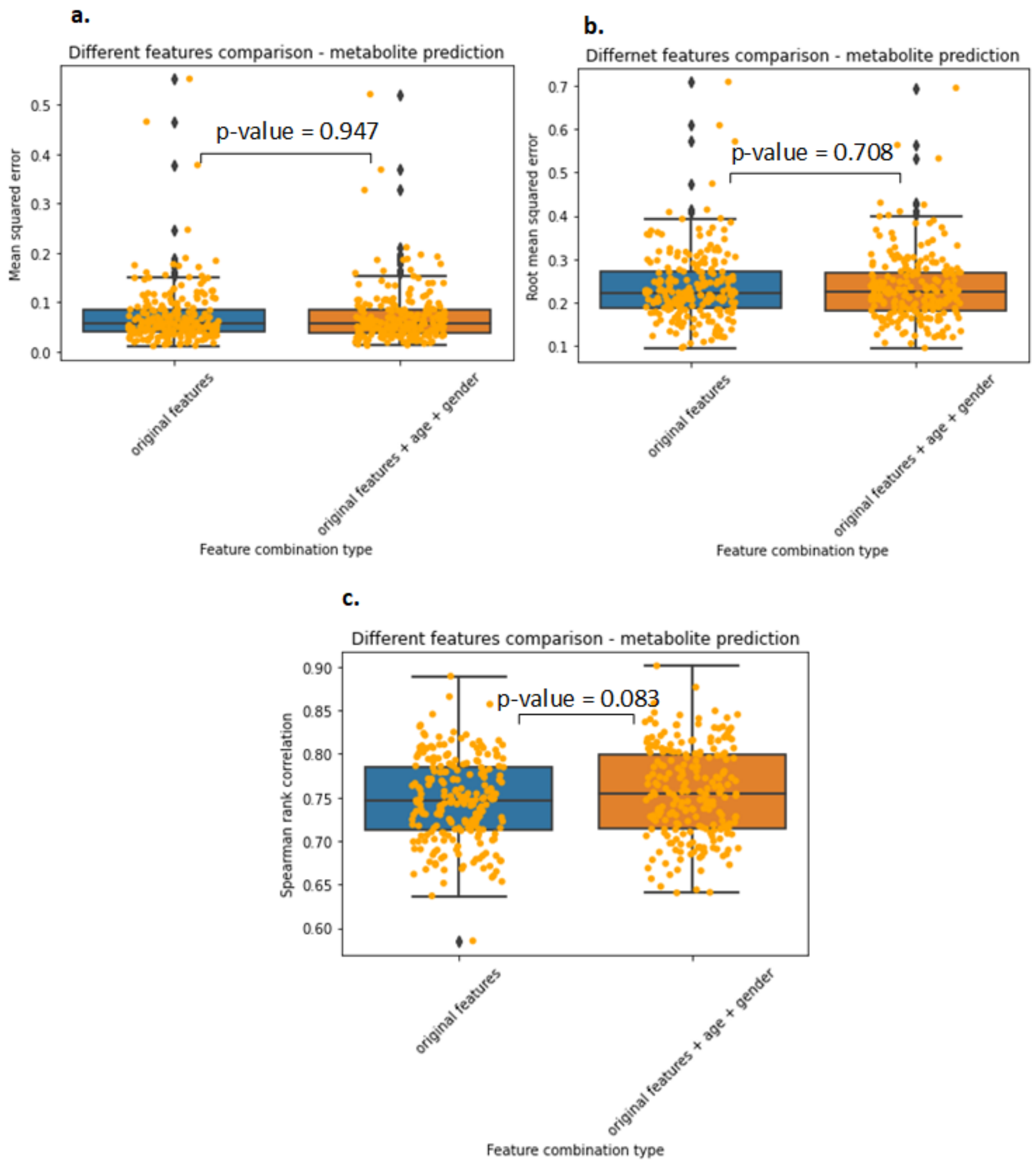


**Figure S4. Different feature set combination comparisons in all metabolite models based on the average of five-fold cross validation results. a-c** The boxplots of (**a**) MSE, (**b**) RMSE, (**c**) Spearman rank correlation.

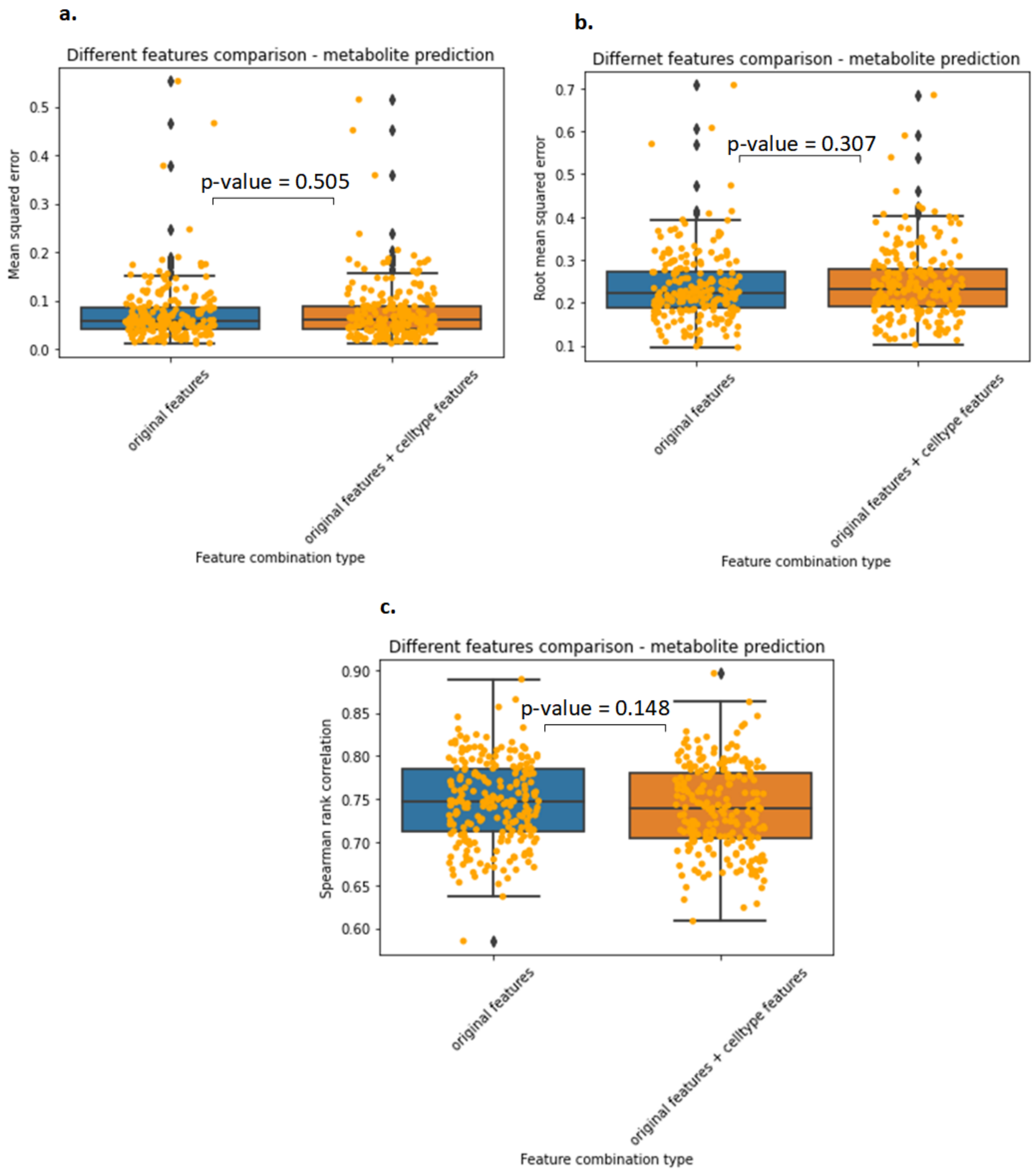


**Figure S5. Different feature set combination comparisons in all metabolite models based on the average of five-fold cross validation results. a-c** The boxplots of (**a**) MSE, (**b**) RMSE, (**c**) Spearman rank correlation.

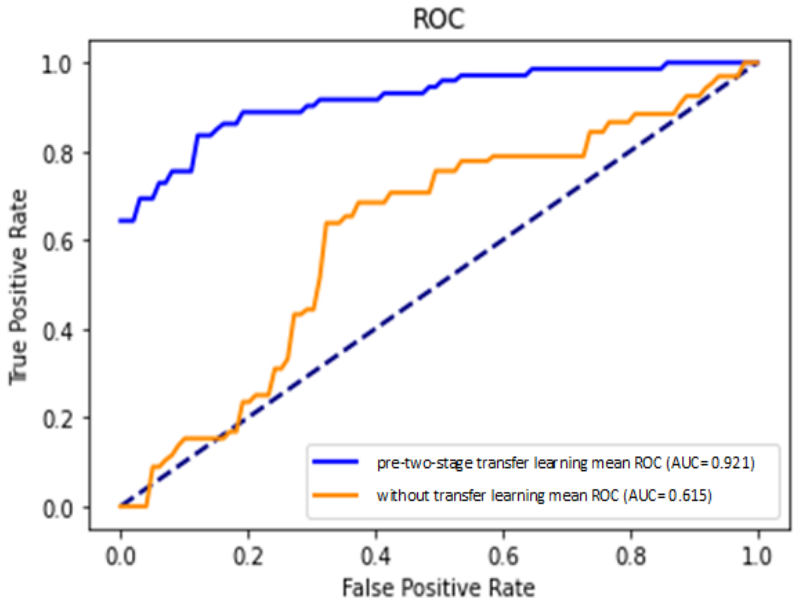


**Figure S6. Receiver operating characteristic curve for clinical prediction framework with pre-two-stage transfer learning and without transfer learning.**

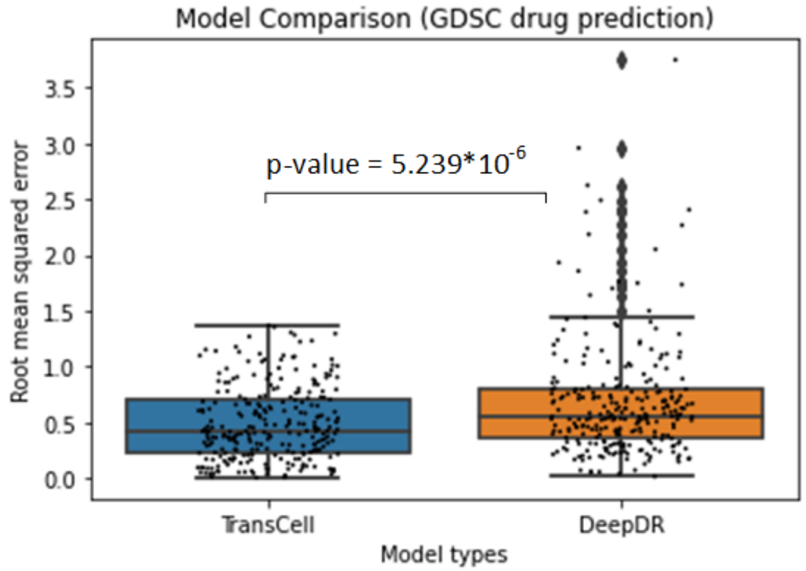


**Figure S7. Model Comparisons between TransCell and DeepDR for cancer cell line (CCLE) drug response prediction based on the genomics of drug sensitivity in cancer (GDSC) project.** The boxplots show the root mean squared error in log-scale $IC_{50}$ for TransCell and DeepDR, respectively.

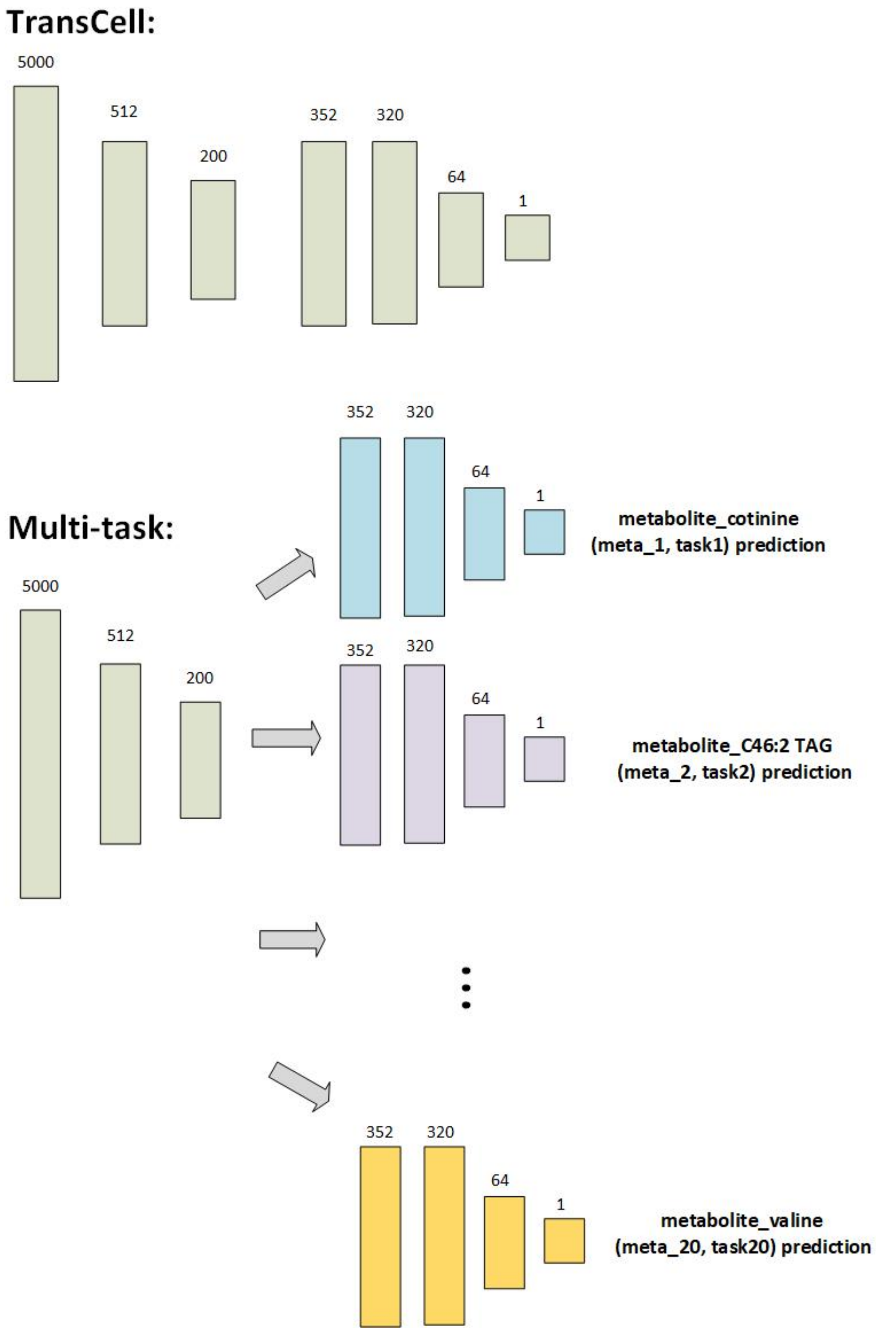


**Figure S8.** The architecture of multi-task learning model for metabolite predictions.

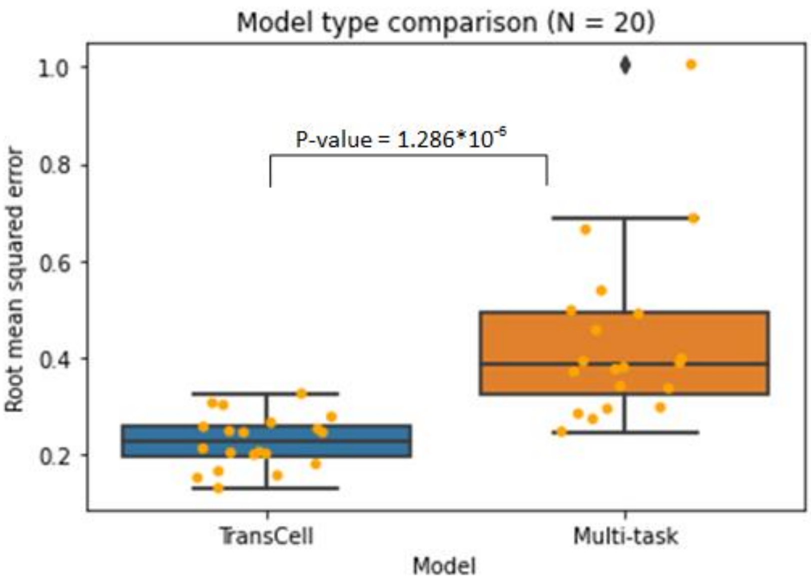


**Figure S9.** The boxplot of root mean squared error between the same 20 metabolite predictions for model type comparison.
